# Resolving Non-identifiability Mitigates Bias in Models of Neural Tuning and Functional Coupling

**DOI:** 10.1101/2023.07.11.548615

**Authors:** Pratik Sachdeva, Ji Hyun Bak, Jesse Livezey, Christoph Kirst, Loren Frank, Sharmodeep Bhattacharyya, Kristofer E. Bouchard

## Abstract

In the brain, all neurons are driven by the activity of other neurons, some of which maybe simultaneously recorded, but most are not. As such, models of neuronal activity need to account for simultaneously recorded neurons and the influences of unmeasured neurons. This can be done through inclusion of model terms for observed external variables (e.g., tuning to stimuli) as well as terms for latent sources of variability. Determining the influence of groups of neurons on each other relative to other influences is important to understand brain functioning. The parameters of statistical models fit to data are commonly used to gain insight into the relative importance of those influences. Scientific interpretation of models hinge upon unbiased parameter estimates. However, evaluation of biased inference is rarely performed and sources of bias are poorly understood. Through extensive numerical study and analytic calculation, we show that common inference procedures and models are typically biased. We demonstrate that accurate parameter selection before estimation resolves model non-identifiability and mitigates bias. In diverse neurophysiology data sets, we found that contributions of coupling to other neurons are often overestimated while tuning to exogenous variables are underestimated in common methods. We explain heterogeneity in observed biases across data sets in terms of data statistics. Finally, counter to common intuition, we found that model non-identifiability contributes to bias, not variance, making it a particularly insidious form of statistical error. Together, our results identify the causes of statistical biases in common models of neural data, provide inference procedures to mitigate that bias, and reveal and explain the impact of those biases in diverse neural data sets.

**Author Summary:** Experimental data of interacting cells under the influence of external as well as unobserved factors are ubiquitous. Parametric models are often used to gain understanding of the processes that generated such data. As such, biological understanding hinges upon accurate inference of model parameters. Whether and how systemic parameter bias manifests in such models is poorly understood. We study this issue in the specific context of estimating the static and dynamic interactions of simultaneously recorded neurons influenced by stimuli and unobserved neurons. Through extensive numerical study and analytic calculations, we identify and mitigate bias in such models. When applied to diverse neural data sets, we found that common models and inference procedures often overestimate the importance of coupling and underestimate tuning. In contrast to common intuition, we find that model non-identifiability contributes to estimation bias, not variance, making it a particularly insidious form of statistical error. As the experimental and statistical issues examined here are common, the insights and solutions we developed will likely impact many fields of biology.

## 2 Introduction

Sensation, cognition, and action are produced by the interaction of neurons distributed throughout the brain. In the central nervous system, all neurons are driven by the activity of other neurons. New technologies are enabling simultaneous recording from increasingly large numbers of sensors distributed across brain areas. For example, the activity of multiple single-units in the primary visual cortex (PVC) of monkeys can be recorded in response to a visual presentation of oriented gratings (Fig. 1a-b); microelectocorticography (*μ*ECoG) from rat auditory cortex can record responses to pure tone sounds (Fig. 1d-e), and multiple single-units can be recorded in the hippocampus of a rat navigating a maze (Fig. 1g-h). These data open the door to models that capture the influences of neurons on each other (Fig. 1b,e,h). At the same time, we are still only able to measure a tiny fraction of the neurons in the brain. As a result, models that seek to explain the activity of one neuron based on the activity of other, simultaneously recorded cells, need to account for the influences of all of the unmeasured neurons. One approach to doing so is to summarize the activity of those unmeasured neurons based on things we can measure, including external factors such as stimulus parameters (Fig. 1a,d,g). The result is a model that seeks to explain neural activity in terms of the influence of other, measured neurons, and the influence of stimulus attributes (i.e., tuning). Another way that unobserved neurons can contribute to neural activity is to create correlations across recorded neurons in the fluctuations about the mean response to the same stimulus (i.e., ‘noise correlations’,Fig. 1c,f,i), which may be described as an unobserved influence. Thus, there are multiple influences that can be used to model neural activity. Determining the influence (i.e., couplings) of groups of single neurons on each other, and the magnitude of those influences relative to the influence of e.g., external stimuli, is important to understanding brain functioning. Modern data sets with many simultaneously recorded neurons bring with them the opportunity to gain deeper insight into the brain computations underlying diverse functions, as well as the challenge of extracting that insight from the data itself.

**Figure 1:**
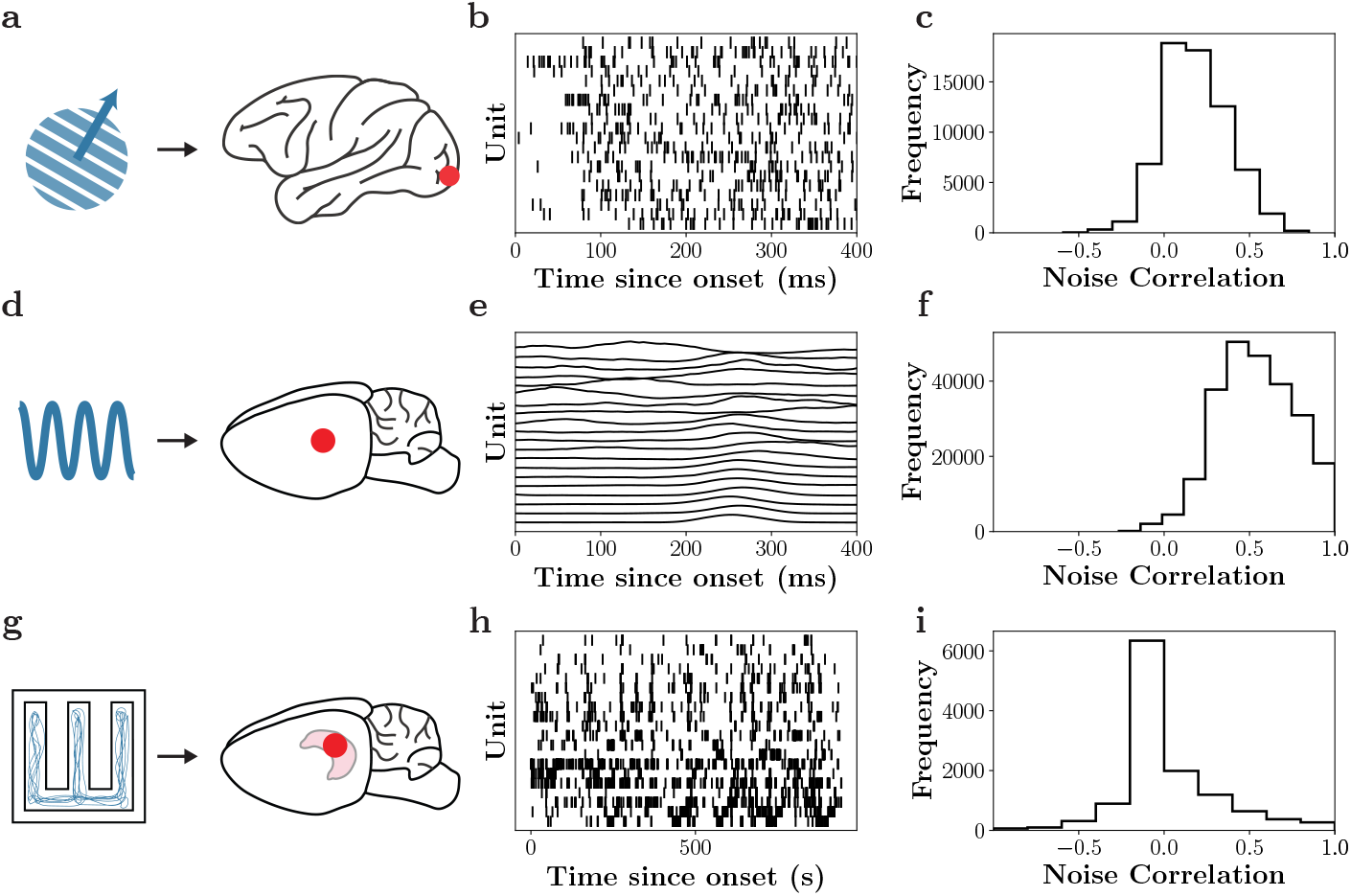
Neural activity depends on external stimuli, simultaneously recorded neurons, and unobserved neural activity. Each row corresponds to a separate neuroscience dataset. **a-c**. Single-unit spikes recorded from macaque monkey primary visual cortex (red dot on the right) in response to drifting gratings (blue diagram on the left). **b**. Spike rasters for 20 distinct single units as a function of time from the onset of stimulus. **c**. The distribution of pairwise noise correlations across the neural population, calculated for each pair of units. **d**. Micro-electrocorticography (*μ*-ECoG) recordings (*z*-scored H*γ* response) from rat primary auditory cortex (right) in response to tone pips at varying frequencies (left). **e**. The high-*γ* response for 20 different electrodes as a function of time from stimulus onset. **f**. The distribution of pairwise noise correlations for the *μ*-ECoG dataset. **g**. Single-unit spikes recorded from rat hippocampus (right) during a spatial decision-making task in a maze (left). **h**. Spike raster from 20 single units as a function of time in the maze. **i**. The distribution of pairwise noise correlations across the neural population, calculated for each pair of units.

Statistical models provide a powerful set of tools for estimating the influence of multiple factors on neural activity [1, 2]. In particular, parametric models, such as generalized linear models, are appealing because the values of estimated parameters specify which factors are important and how important they are in predicting neural activity [3–10]. In systems neuroscience, there are a handful of common parametric modeling approaches. For example, tuning models describe how the activity of individual e.g, neurons are modulated by an external variable (e.g., sensory stimuli), but neglect to consider how other recorded neurons impact the modeled neuron; likewise coupling models describe how the activity of neurons is modulated by other simultaneously recorded neurons, but neglect to capture dependence on external variables. Understanding what factors (i.e., variables) are omitted by these models aids in constructing more complete models. For example, previous work has examined how the inclusion of functional coupling in a tuning model modulates the estimated influence of tuning on single neuron activity [5, 6, 8]. When a coupling and tuning model was fit to data, the ensuing modulation of neural activity by tuning was observed to be greatly reduced compared to a tuning-only model [8]. This has been interpreted as demonstrating that the activity of individual neurons, while modulated by external variables, is primarily driven by the activity of other simultaneously recorded neurons. From a statistical perspective, the explanation is that much of the response modulation that was being attributed to the external variable (tuning) was being explained away by the inclusion of the internal variables (coupling), which were omitted in the tuning only model (i.e., they were omitted variables). While this is certainly intuitive and potentially true, whether and how bias in model estimation affects these results is unknown. Bias in a parametric model is a systematic deviation of an estimated parameter from its ground-truth value. In order to interpret inferred parameters it is crucial to understand the existence of biases [11]. To the extent possible, these biases should be reduced. However, our understanding of the causes of biased inference is nascent, and hence methods to reduce those biases are lacking. As such, our ability to interpret models suffers, potentially impacting conclusions.

The potential for systematic bias arising from omitted variables, a particular type of model structure misspecification, has been studied when estimating functional coupling [8]. Unobserved neural activity has been specifically identified as one potential roadblock to building unbiased models of recorded neural activity [12, 13]. Some neural datasets have been shown to be better predicted from an unobserved shared latent variable and self-history compared to the history of all simultaneously observed neurons [14]. Incorporating latent variables to represent unobserved movements or neural activity into an encoding model on a reaching dataset was shown to improve decoding of the hand [15]. These results highlight the importance of considering the impact of unobserved neurons in models of population neural activity through the inclusion of latent variables. However, little work has been done to understand whether modeling unobserved neural activity via latent variables can mitigate systematic biases (e.g., omitted variables bias). That is, even in a model that has no omitted variables, can inference in that model be biased? In general, it is poorly understood if and how biased parameter estimates can result from the interaction of the structural form of a model and the inference algorithms used to estimate model parameters. To reiterate, if the model does not capture the structure in the data or if the inference is inaccurate, the resulting parameter estimates may be sufficiently biased to jeopardize scientific conclusions.

Here, we systematically address these issues through numerical simulations, analytic calculations, and application to diverse neural data sets. Throughout, we consider how inference interacts with model structure to affect bias in parameter estimation. To gain intuition, we begin with a static model, where all parameters are time-independent. In synthetic data from this model, we show that accurate variable selection in coupling and tuning models with latent variables can eliminate biased parameter estimates.Application and comparison of the models and inference procedures in two real neural data indicates that the parameter biases are substantial. Next, we show through both numerical and analytic results that a dynamic model exhibits the same quality of bias as the static model, and again show that bias is mitigated by accurate variable selection. Our numerical and analytic work provides insight into the heterogeneity of results observed in two additional neural data sets. Finally, to gain a deeper statistical understanding of the problem, we return to the static case and prove that structural non-identifiability (i.e., the existence of an infinite family of model parameter values with equal data likelihood) of the coupling-tuning-latent model is formally resolved by specification of model support. In contrast to common intuition, we show that nonidentifiability can contribute to parameter estimation bias, not estimation variance, making it a particularly insidious form of statistical error.

## 3 Results

Our goal was to understand the relative influence of small groups of neurons on themselves, relative to the influence of external variables and latent factors. To that end, we identify potential sources of biases in parametric models of neural data, and develop models and inference procedures to mitigate such biases. We first show that the standard models and inference algorithms of neural coupling and tuning can exhibit bias in synthetic data. The first-order intuition was that biased parameters were a reflection of the omitted variables bias [12, 16], in which estimates of parameters can be biased if a parametric model does not include (i.e., omits) sources of variability in the response variable. Surprisingly, we show that simply formulating a more complete model that incorporates terms for unobserved variables does not mitigate the omitted variables bias. We found that pre-specification of the model support before parameter estimation is required to de-bias parameter estimates. We begin with a description of the models we investigate and an overview of the inference procedures. In the context of static models, we next explore inference issues with numerical simulations, where ground truth is known, and then in two real neuroscience data sets. We next explore inference in dynamical models with numerical simulations and analytic calculations, and then two real neuroscience data sets. We conclude with a consideration of the interaction of structural non-identifiability of the models, its resolution by sparsity, and the impact of non-identifiability on bias in the static case.

### 3.1 Models

We investigated linear Gaussian models that describe how observed neural activity depends on coupling, or the activities of the other neurons in the system (Fig. 2a: dark gray components), and tuning, or the external variables (Fig. 2a: light gray components), as well as the activity of other unobserved neurons (Fig. 2a: light blue components). For simplicity, we refer to the observed neural population as being composed of “neurons”, although units of a neural recording do not always correspond to neurons (e.g., electrodes in an ECoG array). In agreement with many experimental findings, we assume some amount of sparsity in both the tuning and coupling parameters (this will turn out to be a critical assumption for inference). Suppose that the data measured the activity of a population of *N* neurons during an external variable characterized by *M* features. We denote the neural activity using an *N* -dimensional vector **y**, and the external variables using an *M* -dimensional vector **x**. We consider both static (i.e., time-instantaneous) and dynamic (i.e., time-dependent) models of the relationships between neuronal and external variables. In the static case, we model the activity of a target neuron, *y*_*i*_, separately from the non-target neurons, **y**_¬*i*_. For notational convenience, we choose a convention where **y**_¬*i*_ is an *N* -dimensional vector, such that there are *N* + 1 neurons in the observed population. In the dynamic case, the neural activity and the external variable at each time *t* can be represented by *N* - and *M* -dimensional vectors **y**_*t*_ and **x**_*t*_, respectively. There is no distinction between target and non-target neurons in the dynamical case. Throughout the text, we will use bold letters for vector variables.

**Figure 2:**
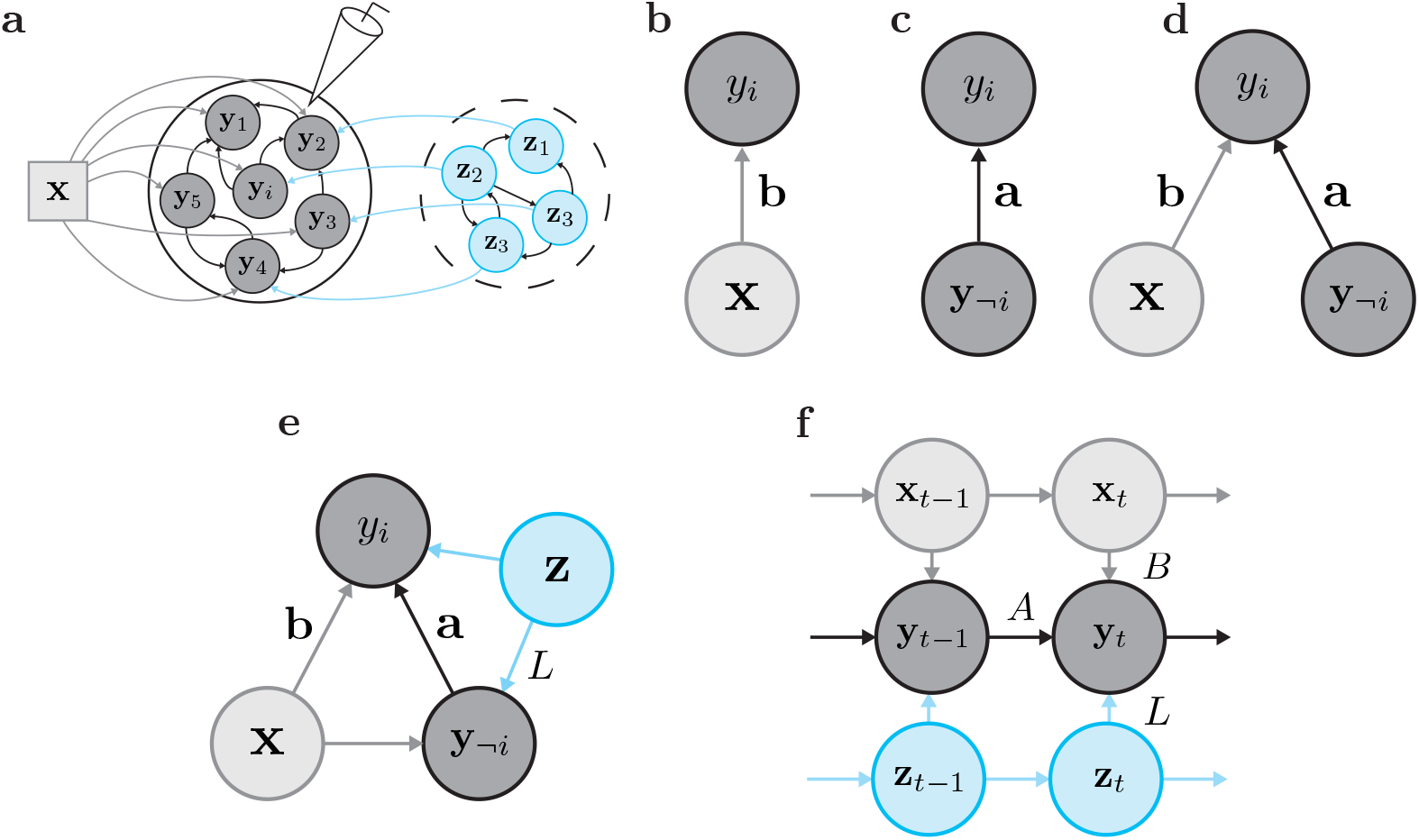
Models of neural activity capturing the impact of tuning, functional coupling, and unobserved influences. **a**. Neural datasets are comprised of recordings (electrode) from observed neurons **y** (center circle) that respond to an external stimulus **x**. The recording apparatus may fail to capture unobserved activity **z** (dashed circle). The recorded neuronal activity depends on the external stimulus (tuning), the within-population interactions (functional coupling) and the unobserved activity. **b-d**. Commonly used systems neuroscience models capturing tuning and functional coupling include the **b**. tuning model, where neuronal dependence on the external stimulus is modeled; **c**. functional coupling model, where a neuron’s dependence on neighboring neurons is modeled; and **d**. the coupling and tuning (CoTu) model, which models both factors simultaneously. **e**. The static CoTuLa model extends the tuning and coupling model by simultaneously capturing the joint impact of external tuning **x** and unobserved activity **z** on the target neuron *y*_*i*_ and non-target neurons **y**_¬*i*_. In the graphical model, **z** is a latent variable (light blue coloring). **f**. A dynamic extension of the static CoTuLa model, which captures the external stimuli and latent variable influence on the neural population at each time point. The model also incorporates both temporal correlations in the stimulus and the latent variable.

#### Static CoTu model

Tuning models (Fig. 2b) and coupling models (Fig. 2c) specify how a target neuron *y*_*i*_ depends on the external variable **x** or the non-target neurons **y**_¬*i*_, respectively. Coupling-Tuning (CoTu) models simultaneously describe a target neuron’s dependence on external variables and non-target neural activity. In the linear-Gaussian setting, a CoTu model can be formulated as:

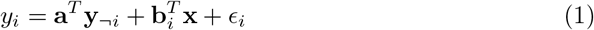

where **a** is a vector of *N* coupling parameters, **b**_*i*_ is a vector of *M* tuning parameters. Finally, *ϵ*_*i*_ is a noise term, assumed to be drawn from a zero-mean Gaussian distribution. **Static CoTuLa model**. The CoTu model omits the effect of any unobserved neurons on the observed population (Fig. 2a: light blue population). To account for unobserved activity, we formulate a static Coupling-Tuning-Latent (CoTuLa) model, as depicted in Fig. 2e. In the CoTuLa model, the target neuron depends jointly on the external variable (e.g., stimulus), non-target neurons, and an unobserved variable. We introduce a *K*-dimensional latent state vector **z**, which acts as a low-dimensional representation of the unobserved neurons. The model additionally incorporates non-target neuron dependence on the stimulus and unobserved activity, more accurately reflecting the neural data generation process to the extent possible within a static framework.

The static CoTuLa model capturing the generation of neural activity at the trial level is:

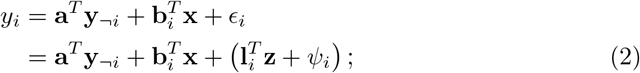

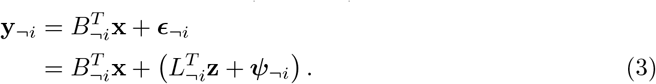

Note that, in Eqn. Eq. 2 and Eq. 3, the target neuron has a special status, in that, Eqn. Eq. 3 does not have neuronal signals on the right-hand-side of the equation. Similar to a Factor Analysis model, the latent state generates variability in the model via a lowdimensional shared component, defined by *L*^*T*^ **z** = [*L*_¬*i*_, **l**_*i*_]^*T*^ **z** and a private component **Ψ** = (***ψ***_¬*i*_, *ψ*_*i*_) which is drawn from a zero-mean Gaussian with learned variance. Note that 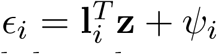 and 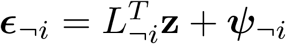 are now used to refer to the total unobserved variability that cannot be captured by the observed parameters.

#### Dynamical CoTu model

We next formulated a dynamical model of neural activity. As in the static model, the dynamic model describes how neural activity depends on coupling to other neurons in the system, as well as tuning to external signals. Suppose that the data simultaneously measured the activity of *N* neurons sampled at *T* time points. This can be summarized in a time series *Y* = {**y**_1_, …, **y**_*t*_, …, **y**_*T*_ }, where the *N* -dimensional vector **y**_*t*_ represents the neural activity at time *t*. At the same time, there are also *M* features of the measured external variable, summarized in a time series *X* = {**x**_1_, …, **x**_*t*_, …, **x**_*T*_ }, where each **x**_*t*_ is an *M* -dimensional vector. Specifically, this work focuses on a 1-step vector autoregressive (VAR_1_) model with an additional tuning term. We write down the dynamical Coupling-Tuning (CoTu) model as:

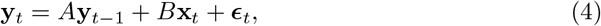

where *A* is an *N* × *N* coupling matrix that captures the dynamic functional coupling between the observed neurons, and *B* is a *N* × *M* tuning matrix that summarizes how the neurons are influenced by the external input. Note that unlike in the formulation of the static model, there is no distinction between the target neuron and the non-target neurons.

#### Dynamical CoTuLa model

As in the static case, we explicitly model spatiotemporal correlations in the unobserved variability by assuming a decomposition ***ϵ***_*t*_ = *L***z**_*t*_ + ***ψ***_*t*_, where **z**_*t*_ is itself a latent dynamical variable, and *L* is an *N* × *K* latent coupling matrix that describes how **z**_*t*_ influences each neuron’s activity. The additional private noise ***ψ***_*t*_ ∼ (**0**, Σ) is assumed to be an *N* -dimensional independently and identically distributed (i.i.d.) Gaussian variable. This allows us to write the dynamical Coupling-Tuning-Latent (CoTuLa) model (see the graphical model in Fig. 2f) as:

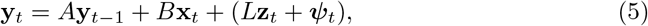

We modeled the dynamics of *Z* = {**z**_*t*_} using a separate VAR_1_ process:

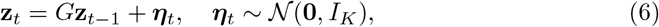

where *K* is the dimensionality of **z**_*t*_, *G* is a *K* × *K* latent coupling matrix, and ***η***_*t*_ is i.i.d. Gaussian noise.

#### Model inference

For both the static and dynamic CoTu models, parameter estimation reduces to a linear regression problem, and ordinary least squares is used. For the CoTuLa models, parameter inference can be conducted via the expectation-maximization (EM) algorithm, because we used linear-Gaussian latent variable models. We developed an EM procedure to estimate the parameters separately for static and dynamical cases; see Methods Section 5.3 and Section 5.4 for details. However, because the model parameters can be sparse, inferring model support is a necessary step in the inference procedure. Inferring model support, a.k.a., variable selection, is the identification of the non-zero parameters in a parametric model. Variable selection in parametric models is commonly achieved through the inclusion of an L1-penalty in the objective function to regularize the parameter values towards zero, often solved with Lasso methods [17]. However, it is known that L1-regularization suffers from both false-positive variable selection as well as bias in parameter estimation in the presence of noise [18].

As we will show, the accuracy of model support can have a dramatic effect on the estimated parameters. We explored three model support regimes: no selection, oracle selection, and inferred selection. With no selection, the values of all parameters are estimated; with oracle selection, the true model supports of the coupling and tuning parameters are provided. We utilized a modular approach to parameter inference, first performing inference of variable selection, followed by estimation of non-zero parameter values, which has been shown to achieve very good results in similar systems neuroscience models [18, 19]. The requirement of accurate sparse model selection in terms of structural non-identifiability of the CoTuLa model is justified in Sections 3.7 and 3.8. A full inference procedure, therefore, requires specifying a method to first conduct selection and a separate method to perform the estimation. In such a modular inference algorithm, different “model supports” or sets of non-zero parameters can be passed into the parameter estimation module (in this paper, either linear regression or EM). This two-stage procedure was done for two reasons: (1) our goal was to disentangle the issues of selection accuracy from estimation bias during inference and show that accurate selection enables unbiased estimation, and, (2) we observed that utilization of L1-regularization during inference in either the CoTu or CoTuLa models did not alleviate the bias we are about to describe (Fig. S.1). For inferred selection in the static models, we used UoI_Lasso_ which infers model supports with near-oracle selection accuracy from data samples (see Methods Section 5.5) [18, 19]. We evaluated the accuracy of the selection procedure, finding that it exhibited moderate to good selection performance in all regimes (Fig. S.2). The procedure tended to suffer from false positives, implying that the procedure may indicate a feature was predictive of the target neuron when in fact, it was not. In the context of CoTu and CoTuLa inference, however, false positives are less detrimental than false negatives, as the latter can induce additional omitted variables bias [12, 20]. Furthermore, false positives can be mitigated if the corresponding estimated parameter value is close to 0, having negligible impacts on prediction. To infer model supports for the dynamic models, we designed and used a normalization-and-cutoff method based on the CoTu fit, which captures the correlated structure of OLS errors in the case of small *K* (see Methods Section 5.5 for details). This method of variable selection worked very well in practice in the synthetic data. In the following sections, we will use the terms “sparse inference in the CoTu model” or “sparse inference in the CoTuLa model” to refer to the modularized inference procedures that use the parameter estimation method developed for the CoTu or CoTuLa models, respectively. See Methods Section 5.5 for more details on the selection procedures.

### 3.2 Unbiased parameter estimation requires accurate model selection during inference of the CoTuLa model

As described above, the CoTuLa model more accurately reflects the neural data generation process than the CoTu model. We therefore evaluated the presence of biased inference in those models on synthetic data generated from the CoTuLa model. We performed a systematic, large-scale numerical experiment with two hyperparameters of interest: the mean of the coupling parameters and the mean of the tuning parameters. For each hyperparameter, we examined 5 values in [− 1, 1] (for a total sweep of 5 × 5 = 25 hyperparameter configurations). For simplicity, we structured the unobserved activity to reproduce uniform noise correlations equal to 0.10. For each hyperparameter pair, we generated 10 models with different sets of parameters. For each model, we generated 30 datasets and performed inference over 3 folds of the data. We estimated statistics by taking averages across datasets and folds, followed by a median across models, and finally a median across parameters within a model. We calculated the biases for the ground truth non-zero coupling and tuning parameters as a function of the underlying hyperparameters. See Methods Section 5.2 for further details.

We focused on examining the coupling and tuning parameter estimates for the target neuron (**a** and **b**), since these parameters are shared between the CoTu and CoTuLa models. We quantified the accuracy of the coupling and tuning parameter estimates obtained by fitting a CoTuLa model with the inference procedures described in Methods Section 5.3. We compared these estimates to those provided by CoTu inference on the same synthetic data. This allowed us to both assess the impact of model support in CoTuLa inference, as well as characterize the manner in which CoTu inference may be biased when the data contains unobserved sources of variability. As outlined in the previous section, we divided the overall inference procedure into two stages: selection, where the non-zero coupling and tuning parameters are identified, and estimation, where the values of those non-zero parameters are determined. In the current analysis, we considered three selection regimes: oracle selection, inferred selection, and no selection. In oracle selection, each estimation was performed on the true model support of the CoTuLa model. However, oracle selection is not available for real data. As such, for inferred selection, we used UoI_Lasso_ which infers model supports with near-oracle selection accuracy from data samples (see Methods Section 5.5) [18, 19]. For no selection, the full parameter set is used for estimation. Expectation-maximization for the CoTuLa model and ordinary least squares for the CoTu model was used. Importantly, both CoTu and CoTuLa inference procedures were provided the same model supports in order to effectively compare how parameter estimates depended on selection accuracy during inference.

In Figure 3, we display the difference between inferred and actual coupling and tuning parameter values (i.e., estimation bias, color bars; color bars are shared for all plots for coupling and tuning). For the three selection sets (oracle selection (Fig. 3a-d), inferred selection (Fig. 3e-h), and no selection (Fig. 3i-l)) the coupling (top row) and tuning bias (bottom row) are plotted as a function of the mean of the tuning (x-axis) and coupling (y-axis) parameter values in the CoTuLa generative models when inference was performed in the CoTu (right column) or CoTuLa models (left column). Across selection sets, we found that inference in the CoTu model suffers from clear biases, which arise from not modeling the unobserved variable (Fig. 3a,c,e,g,i,k). Interestingly, we observed only positive biases for the coupling parameters, while the tuning parameters exhibited negative and positive biases. We further found that, as we might expect, bias magnitude increased with both the tuning or coupling mean (Fig. 3). Furthermore, the tuning parameter bias corresponded to the sign of the underlying distribution: if the distribution is positive, the bias will be negative, and vice versa (Fig. 3). As expected, we observed that tuning biases were qualitatively similar (i.e., tuning modulation is reduced) when the neural response was modeled as a Poisson distribution (Fig. S.3). The effect of the different selection profiles on inference in the CoTu model was minimal.

**Figure 3:**
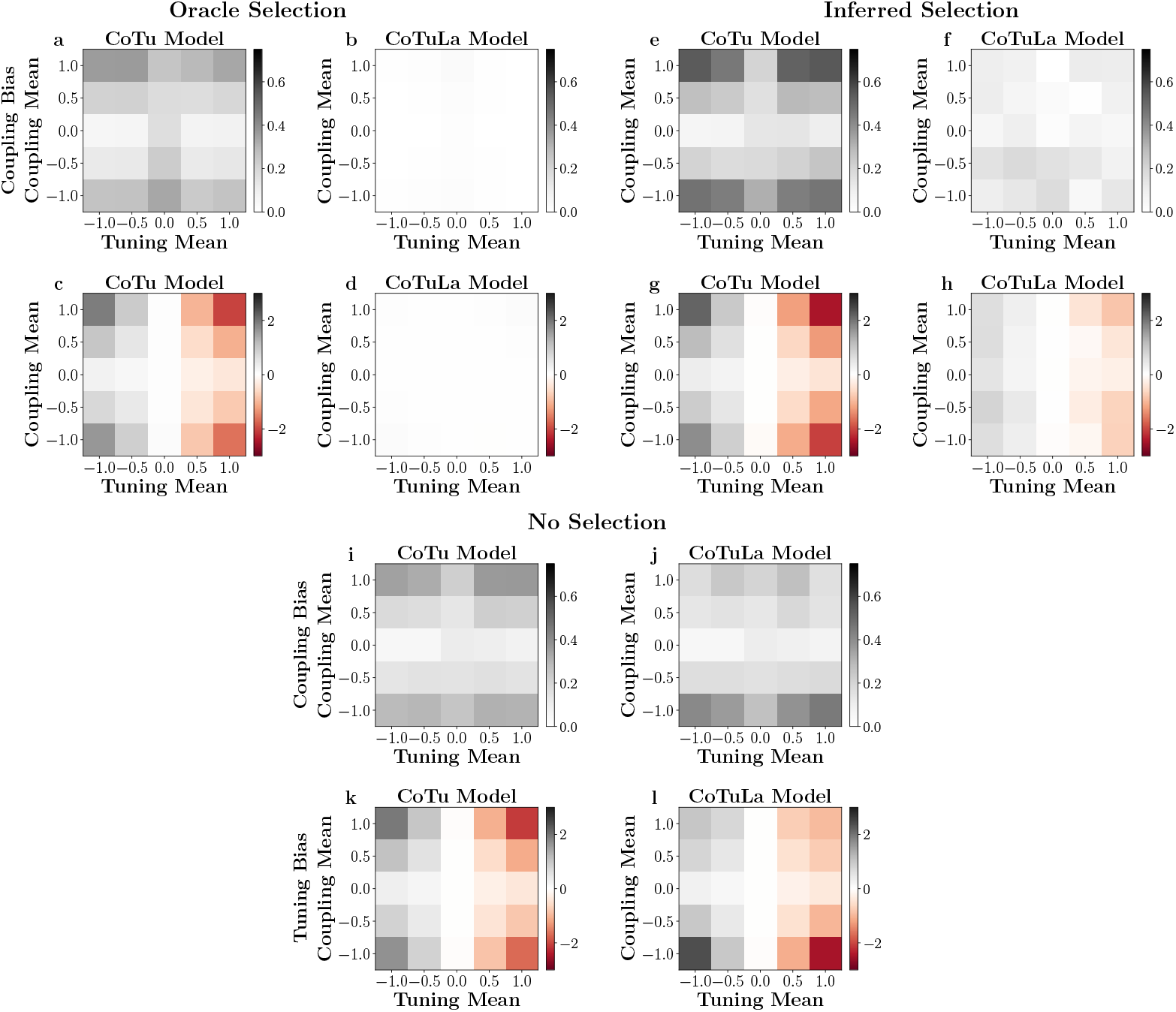
Unbiased parameter estimation requires accurate model selection during inference of the CoTuLa model. Each plot presents normalized bias as a function of the parameter-generating distributions’ means: the tuning mean (*x*-axes) and the coupling mean (*y*-axes). We examined three conditions of model selection: oracle selection (**a-d**.), inferred selection (**e-h**.), and no selection (**i-l**). In each, the top row depicts results for coupling parameters, while the bottom row depicts results for tuning parameters. Tuning and coupling biases are respectively plotted with the same colormaps, shown in the colorbars on the right of each subplot.

In stark contrast, for the CoTuLa model, we found a major effect of the selection set on the magnitude of coupling and tuning parameter bias (Fig. 3b,d,f,h,j,l). In particular, we found that for oracle selection, CoTuLa parameter biases are an order of magnitude smaller (plots were entirely white), indicating what is effectively unbiased estimation. Estimation based on inferred selection for the CoTuLa model resulted in biases that were substantially better than the CoTu model, but slightly worse than for oracle selection (Fig. 3f,h). For no selection, parameter bias in the CoTuLa model was essentially on-par with the CoTu model (Fig. 3j,l). The general pattern of bias dependence on the coupling and tuning means was similar between the CoTu and CoTuLa models. Together, these results demonstrate that inference in the CoTu model is susceptible to biases when unobserved activity exists but is not modeled, and inference in the CoTuLa model mitigates or completely removes those biases, depending on the quality of parameter selection. In the context of real neural data, these results suggest that standard inference in the CoTu model may result in artificially reduced ratios of tuning relative to coupling parameters.

The solution to the linear-Gaussian static CoTuLa model is generally not analytically tractable except in the univariate case, where there is a single tuning parameter, and a single coupling parameter. We analytically calculated the biases obtained by CoTu inference applied to CoTuLa-generated data (Supplementary Information). The biases scale according to the tuning parameters (in agreement with Fig. 3). However, the analytic form of the biases in the linear-Gaussian model are difficult to interpret due to the inherent structural non-identifiability of the model (see Section 3.7).

### 3.3 Sparse inference in the static CoTuLa model mitigates bias in neural data

Previous results comparing coupling and tuning parameters estimated by independent tuning and coupling models with estimates from a coupling-tuning (CoTu) model reported a dramatic decrease in the magnitude of tuning relative to coupling in the CoTu model. This was interpreted as evidence that neuronal activity is primarily driven by the coupling to other neurons, and only weakly modulated by tuning to external variables [6]. However, the results of the previous section demonstrated that this prior result could, in part, reflect bias due to the omission of latent variables combined with poor variable selection. Thus, we next assessed whether sparse inference in the CoTuLa model, which mitigated the biases in synthetic data, had a similar impact on neural data. We considered two datasets: single-unit activity from macaque primary visual cortex (PVC) in response to drifting gratings, and *μ*ECoG recordings from rat auditory cortex in response to tone pips. We fit linear-Gaussian CoTu and CoTuLa models, encoding the tuning parameters using basis functions. We performed inference with the same procedure used on the synthetic data and compared the values of the estimated parameters from fits to the CoTu and CoTuLa models, as well as coupling and tuning models. See Methods Section 5.1 for more details.

The results for the primary visual cortex are presented in Figure 4a-c and for the auditory cortex in Figure 4d-f. Example tuning curves for each data set are presented in (Fig. 4a, d). Here, as elsewhere in this figure, fits from the baseline models (tuning or coupling) are indicated in black, fits from the CoTu model are in grey, while fits from CoTuLa are in red. For both data sets, we observed a substantial reduction in tuning modulation between the Tu and CoTu fits (i.e., grey lines have a reduced min-max range), which was often partially recovered by sparse inference in the CoTuLa models. We quantified the differences between model fits as the minimum-to-maximum distance of each tuning curve. We then directly compared CoTu and CoTuLa tuning modulations for each model fit for each single-unit or *μ*ECoG channel (Fig. 4b, e). In visual cortex data, we observed a general elevation of tuning modulations relative to the CoTu model (points are above the diagonal,Fig. 4b). Across the neural population, the increase in tuning modulation was statistically significant (Fig. 4b: inset: ***: *p* = 7 × 10^−6^, Wilcoxon signed-rank test, *N* = 106). Likewise, in the auditory cortex, the tuning modulations of the CoTuLa model are almost universally elevated relative to the CoTu model (Fig. 4d-e). The CoTuLa tuning modulations were significantly larger than the corresponding CoTu tuning modulations (Fig. 4e: inset: ***: *p* = 2 × 10^−20^, Wilcoxon signed-rank test, *N* = 128). We note that, in both cases, tuning modulations from the CoTuLa model were significantly lower than those from a tuning model alone (Fig. 4b-e: inset, black; b: ***: *p* = 9 × 10^−10^, Wilcoxon signed-rank test, *N* = 106; e: ***: *p* = 2 × 10^−22^, Wilcoxon signed-rank test, *N* = 128), implying that some explaining away of tuning persist with sparse inference in the CoTuLa model.

**Figure 4:**
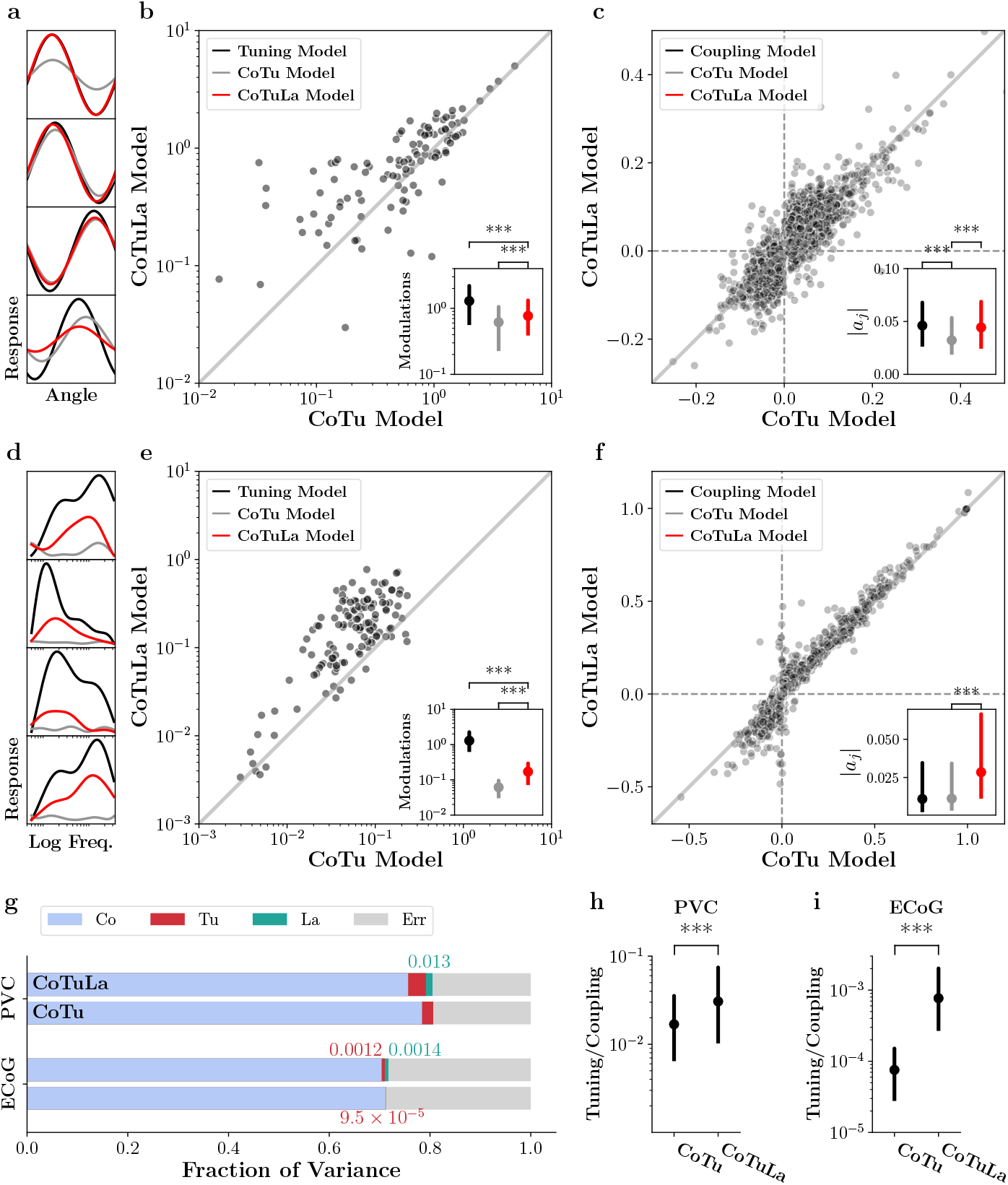
Sparse inference in the static CoTuLa model mitigates bias in neural data. **a-c**. Results from monkey V1 (PVC) data. **a**. Example fitted tuning curves on single units using a tuning model (black), CoTu model (gray) and CoTuLa model (red). **b**. Tuning modulations obtained from CoTu model (*x*-axis) compared to those obtained from CoTuLa model (*y*-axis) across the single-units in the population. Inset depicts the distribution (median and IQR) of tuning modulations, for the tuning (black), CoTu (grey), and CoTuLa (red) models. Note the log-scale on the axes. **c**. Comparison of coupling parameters between the CoTu (*x*-axis) and CoTuLa (*y*-axis) models. Points denote coupling parameters across models. Inset depicts the distribution (median and IQR) of coupling parameter magnitudes, for the tuning (black), CoTu (grey), and CoTuLa (red) models. **d-f**. Results on rat *μ*-ECoG data, with similar plots as the top row. **c**. Example fitted tuning curves on single electrodes **d-f**. Same as **a-c**. but for rat primary auditory cortex data. **d-e**. Comparison of tuning modulations. **e**. Comparison of coupling parameters. **g**. Fraction of variance in the neural responses captured by each term in the CoTu and CoTuLa models, shown in a stacked bar plot. **h**. Tuning-coupling ratios, calculated from the variance fractions, based on the CoTu and CoTuLa model inference procedures. Data are displayed as medians and IQR (interquartile range) over all neurons in the fit. Significance markers denote *p <* 10^−3^ for Wilcoxon signed-rank test.

We next compared the coupling coefficients fitted from the CoTuLa model to the corresponding coupling coefficients in the CoTu model on a per-unit basis (Fig. 4c,f). Interestingly, we observed diverse effects, with CoTuLa parameter estimates exceeding, equaling, or below their CoTu counterparts (Fig. 4c, f). We compared the magnitude of the coupling parameter values at the population level, finding that CoTuLa coupling parameter magnitudes were significantly higher than their CoTu counterparts (Fig. 4c,f, insets, c: ***: *p* = 4.3 × 10^−81^, Wilcoxon signed-rank test, *N* = 106; e: *p* = 1 × 10^−100^, Wilcoxon signed-rank test, *N* = 128). These results contrast with expectations from the synthetic results (Fig. 3), where we found coupling parameters to be overestimated by the CoTu model (thus, we might expect to see points lie below the identity line in Fig. 4). The observed heterogeneity in the impact of CoTuLa model inference, which may stem from the diversity of noise correlations in Fig. 1c,f, where both positive and negative noise correlations are observed in the data (for simplicity, only positive noise correlations were utilized in the numerical results of the previous section). On the whole, these results demonstrate that sparse inference in the static CoTuLa model mitigates the omitted variables bias in neural data, which removes some of the explaining away effect in the CoTu model, but not all of it.

We summarized the contributions of the coupling, tuning, and latent terms in the CoTuLa model to the target neuron’s activity. We calculated the fraction of variance that each term contributed to the target neuron’s response across trials and stimuli (Fig. 4g). We observed that, in both PVC and ECoG datasets, the dominant contribution to neural activity modulation came from coupling (blue bars). A much smaller contribution came from the tuning terms, particularly in the ECoG dataset (green bars). The latent term, meanwhile, was even smaller in the PVC data, but roughly equal to the tuning contribution in the ECoG dataset (red bars). Finally, we directly compared the contributions of tuning relative to coupling estimated from the CoTu and CoTuLa model in (Fig. 4h-i). We found that the ratio of tuning to coupling was significantly higher for estimates from the CoTuLa model compared to the CoTu model (Fig. 4h: *p* = 2 × 10^−6^, Wilcoxon signed-rank test, *N* = 106; i: *p* = 6 × 10^−21^, Wilcoxon signedrank test, *N* = 128). Together, these results demonstrate that sparse inference in the static CoTuLa model significantly increases the magnitude of tuning modulation relative to coupling compared to the CoTu in neural data, with the primary effect coming from an increase in tuning modulation. This indicates that inference in the CoTu model likely suffers from biased parameter estimates in these data sets.

### 3.4 Temporal correlations in unobserved variability can create biases in the dynamical CoTuLa model

Up to this point, we have considered a static model of functional coupling and tuning with and without latent variables, and how inference in these models may suffer from bias depending on the accuracy of model selection. The initial focus on a static model was, in part, motivated by the fact many classical neurophysiology studies deployed an experimental design well matched to this model. For example, in both of the experimental data sets examined, the external stimuli were randomly presented in a trialized manner with long inter-stimulus intervals, in large part to enable an analysis of responses without considering dynamics. However, natural biological phenomena, including ethological behaviors and stimuli, as well as brain activity, has non-trivial temporal correlations. Indeed, many modern neurophysiology experiments, especially ones with a behavioral component (e.g., rats navigating a maze), focus directly on the analysis of the relationship between neural dynamics and dynamics of the external world. Is it possible that parameter bias is resolved by considering a dynamical model, where the time-varying values of each variable are taken into account with explicit time indices (Fig. 2f), instead of being collapsed into a single distribution (Fig. 2e)? Here we show that bias of coupling estimation persists in dynamical models when there is temporal correlation in the unobserved variability (i.e., noise) (Eq. 5), as is the case in neurophysiology data [21, 22]. Furthermore, when temporally correlated noise is combined with temporal correlations in the external variables, we find that tuning parameter estimation becomes biased. Even when both neural activity and external variables are measured at specific times, temporal correlation in the generating sources can create “simultaneity” in the model, as diagrammed in the quadrangular graphs in Fig. 5a (analogous to the triangular graphs in the static CoTuLa model). Importantly, the simultaneity in this context only means that there are more than one sources of variability that co-influence the dynamical variables corresponding to the observed neural activity (Fig. 5a, black circles), regardless of which point in time the variability originated. As we will see, when using model parameters similar to the static case (e.g., positive correlations, strictly positive tuning parameters, etc.) the nature of the bias in the dynamic case is qualitatively similar to the static case.

**Figure 5:**
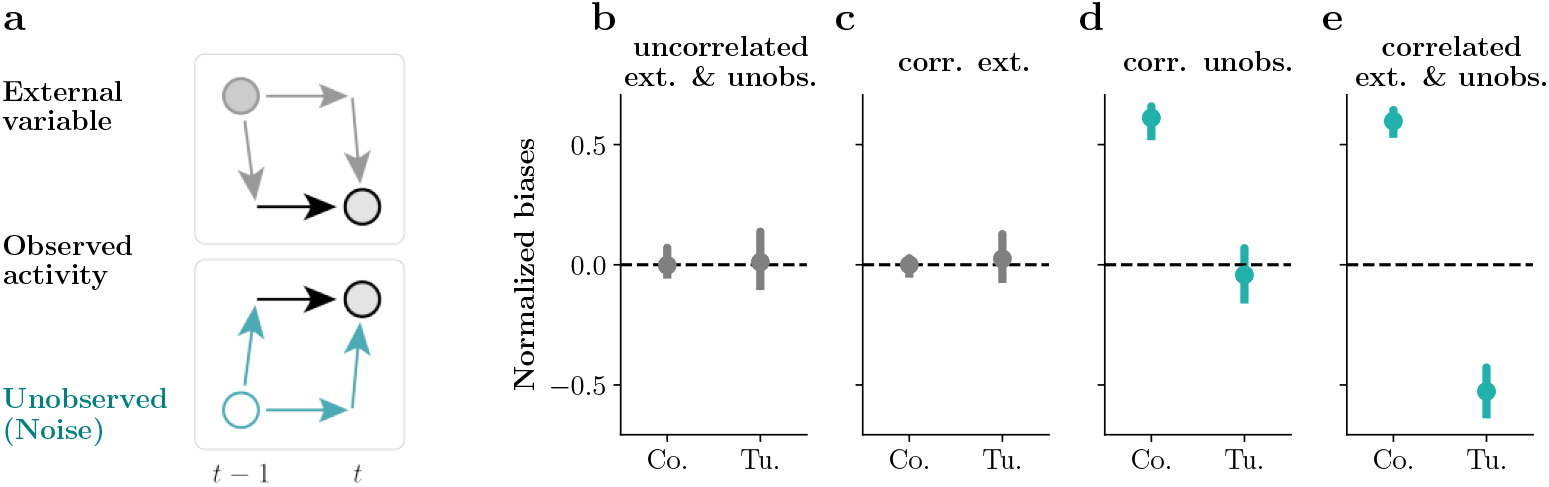
Temporal correlations in unobserved variability can create biases in the dynamical CoTuLa model. Each panel visualizes the median and interquartile range of the normalized biases for the dynamical functional coupling (Co) and tuning (Tu) parameters of the model, with 50 realizations each. **a**. Schematic illustration of the origin of simultaneity in the dynamical model, in the presence of temporal correlation. **b-e**. Systematic biases in OLS estimates. Data were generated, according to four different conditions of the CoTuLa model: **b**. no temporal correlation in either stimuli or noise, **c**. temporal correlation in the stimuli, **d**. temporal correlation in the noise, and **e**. temporal correlation in both stimuli and noise.

To gain analytic insight into potential biases during inference in the dynamic CoTu model, we first derived the biases we would get from naive OLS inference in the dynamic CoTu model (Eq. 4), when the data were generated from the dynamic CoTuLa model (Eq. 5–Eq. 6). For analytic tractability, we considered the univariate case (*N, M, K* = 1) (see Section 5.4.1 for details). We found that the normalized biases for the coupling parameter *a* and the tuning parameter *b* can be written as:

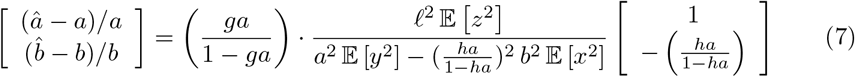

Where *ℓ* is the latent coupling parameter, while *g* and *h* parameterize the strengths of temporal correlation in the latent variable *z*_*t*_ and external variable *x*_*t*_, respectively. In the multivariate case, we numerically evaluated normalized biases in the OLS estimates of the coupling and tuning parameters where the ground truth values are known (see Methods Section 5.2 for details of synthetic data generation, and Methods Section 5.6 for analysis details.)

The numerical results from the multivariate case are consistent with predictions of the analytics in the univariate case. From Eq. 7, we found that the overall normalized biases are scaled by *ga/*(1 − *ga*) and 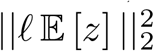 (i.e., the L2-norm of the expected value of the squared unobserved variability). This shows that the OLS estimate has no systematic bias when there is no temporal correlation in the noise (*g* = 0). This analytic prediction from the univariate case was verified by the numerical results in the high-dimensional case. In Figure 5b-e, we plot the normalized biases for the coupling and tuning parameters with and without temporal correlations in the unobserved and external variables. We find that the coupling and tuning biases are centered around zero when there is no correlation in both the unobserved and external variables (Fig. 5b), or when there is temporal correlation in only the external variable (Fig. 5c). Furthermore, the *ℓ*^2^ 𝔼 [*z*]^2^ factor means that the contribution from the unobserved variability should be large in order to create large biases. In the presence of temporally correlated noise, the scaling factor *ga/*(1 − *ga*) is larger when the coupling parameters *a* and *g* are closer to 1; in other words, when there are stronger temporal correlations in both *X* and *Z*. We observed large bias in the coupling term (but not the tuning term) when the unobserved variability had temporal correlations (Fig. 5d).

The tuning bias is further scaled by a factor of −*ha/*(1 − *ha*). With temporally uncorrelated external variables (*h* = 0), the tuning bias vanishes, while the coupling bias remains as long as there is temporal correlation in the noise, *g* ≠ 0, as observed numerically (Fig. 5d). In the presence of temporally correlated external variables, the normalized tuning bias has a negative sign, while the normalized coupling bias has a positive sign (Fig. 5e). These result are consistent with the systematically overestimated coupling parameters and underestimated tuning parameters observed in the static models above.

### 3.5 Sparse inference in the dynamical CoTuLa model mitigates biases in synthetic data

The biases demonstrated in Fig. 5 were introduced by the temporally correlated “noise” due to unobserved influences, and compounded by temporal correlations in the external variables. The nature of the biases in the dynamic CoTu model is similar to the biases observed in the static CoTu model. Therefore, we next tested if sparse inference in the dynamic CoTuLa model mitigates biases in synthetic data.

We took the same set of synthetically generated data as used in the previous section where we demonstrated biases for OLS inference in the dynamic CoTu model, with four different conditions on the generative models of temporal correlation in the external and unobserved variables. For each dataset (observed variables *X* and *Y*), we performed sparse inference in the CoTuLa model to estimate the coupling and tuning matrices *A* and *B*, as well as the latent dynamical variables *Z* and latent coupling matrix *L*. As in the case of static models, we modularized the inference procedure and utilized model support obtained separately from parameter estimation. We contrasted three selection regimes: no selection, inferred selection, and oracle selection. With no selection, the values of all parameters are estimated; with oracle selection, the true model supports of the coupling and tuning parameters are provided.

In Figure 6, we plot normalized biases for coupling and tuning parameters for the CoTu (Fig. 6a-c) and CoTuLa (Fig. 6d-f) models estimated for the three model support regimes and four data generative models. As observed above, estimates based on inference in the CoTu model were systematically biased in the presence of temporal correlations (Fig. 6a). The biases were present and even exacerbated with inferred selections (Fig. 6b), or even with oracle selection (Fig. 6c). When inference in the dynamic CoTuLa model was performed with no selection (Fig. 6d), the estimated coupling and tuning parameters were slightly closer to the true values on average compared to inference in the CoTu model with no selection, but the resulting parameter estimates were more variable. Thus, as with the static case, simply including a latent variable in the model is not sufficient to completely remove biased estimates. Dynamic CoTuLa inference with inferred selection removed the tuning biases in this synthetic example, although not the coupling biases (Fig. 6e). When inference in the dynamic CoTuLa model was performed with oracle selection, however, both the coupling and tuning parameters were inferred accurately and precisely (Fig. 6f). Together with the results of the static model, these results demonstrate that accurate variable selection in generative models can mitigate/resolve bias.

**Figure 6:**
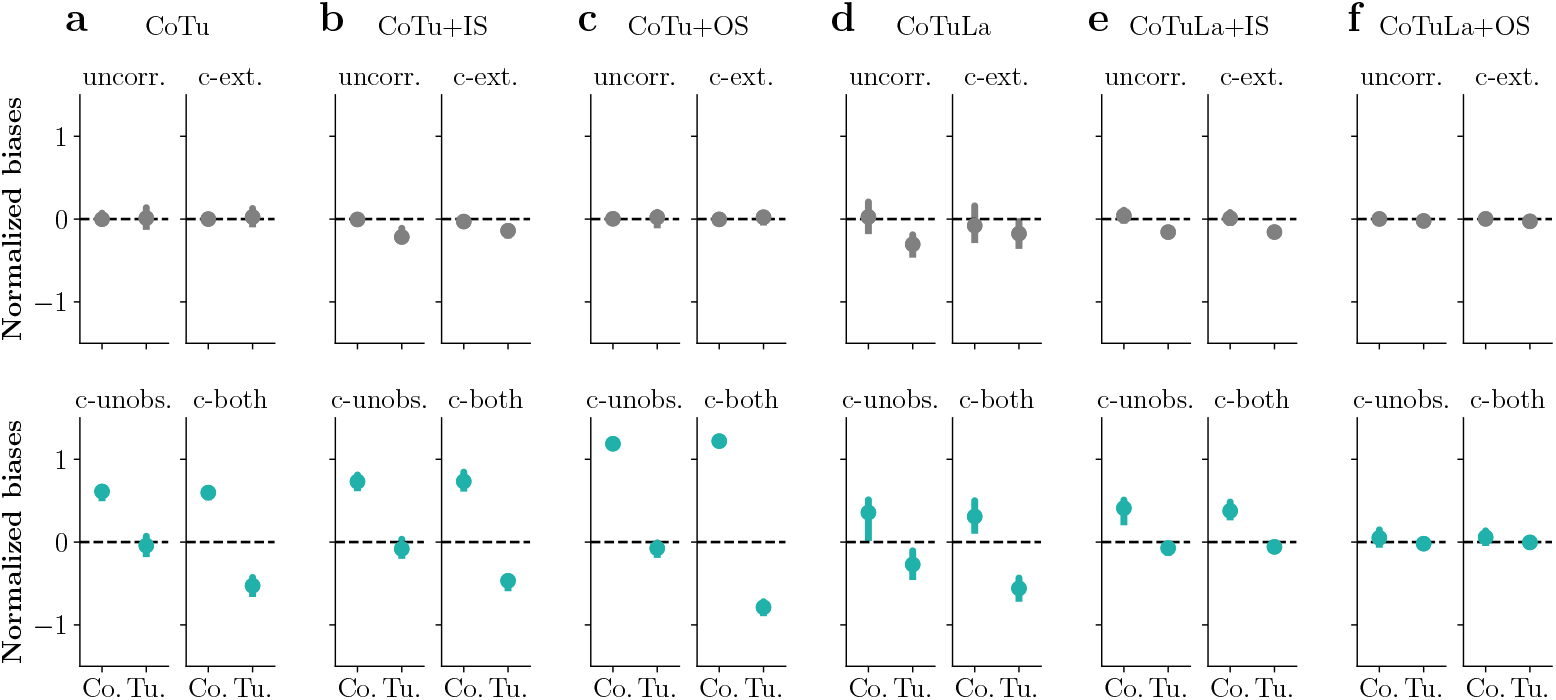
Sparse inference in the dynamical CoTuLa model mitigates biases in synthetic data. Normalized biases for the coupling (Co.) and tuning (Tu.) parameters, under four different generative models with and without temporal correlations. We applied different inference procedures to the same set of data. **a-c**. CoTu model inference, with **a**. no selection, **b**. inferred selection, and **c**. oracle selection. **d-f**. Similar to **a-c**. but for CoTuLa model inference.

### 3.6 Sparse inference in the dynamical CoTuLa model can mitigate bias in neural data

The results above demonstrate that the severity estimation biases, which are introduced by temporal correlations in unobserved variability, is determined by the combination of different terms in the model, not only the latent term (Eq. 7). Specifically, a system with stronger coupling suffers more from biases. Seperately, the strength of temporal correlation in the external variables modulates the tuning biases. In real neural data, it is reasonable to believe that latent variability, reflecting the influence of unobserved neural activity on the recorded population, has non-trivial correlations. Furthermore, across different brain areas, we expect heterogeneity in the strength of the coupling between neurons, while temporal correlations in the external variable (e.g., visual stimulus, location in a maze) will depend on the experimental paradigm. Thus, we next asked: how much of a problem are the omitted variable biases in the analysis of real neural data using dynamical models?

To ascertain the degree to which biased inference impacts the estimation of coupling and tuning in dynamic models, we analyzed two neural datasets that measured neural activity along with time-varying external variables. We examined simultaneously recorded spiking activities from multiple single-units in the macaque primary visual cortex in response to a movie of fast-changing drifting gratings with pseudo-random ordering of orientations [23], and from rat hippocampus CA1 and CA3 during a spatial decision-making task in a maze [24] (see Methods Section 5.1 for details). As we have demonstrated the importance of the accuracy of variable selection to mitigate bias, here, we focused just on the impact of the model used during inference. We performed inference in the dynamical CoTu and CoTuLa models using the same model supports. For the visual cortex data, we divided each data set into four subsets with 30 trials, and fitted each subset. For the hippocampus data, we fitted each experimental session. Units fitted in different subsets were considered distinct units in the following analysis. See Methods Section 5.6 for further details.

We first examined the estimated values of the tuning parameters. The plots in (Fig. 7a,d) display example tuning curves for single-units from PVC and HPC, respectively, estimated from a pure tuning model (black), the CoTu model (grey), and the CoTuLa model (red). In contrast to what was observed for the (different) neural data sets using the static models, these examples suggest negligible differences between the CoTu and CoTuLa models for both the PVC and HPC data sets. We quantified the magnitude of tuning modulation for all single-neurons for the fits from the CoTu and CoTuLa models and plotted them against each other for both data sets (Fig. 7b,e). To explore the potential effects of learning, for the hippocampus data, we visualized the trials that occurred early (purple) and late (black) separately in the scatter plot (Fig. 7e). For both the visual cortex data (Fig. 7b) and the hippocampus data (Fig. 7e), we observed that the inferred tuning parameters are highly consistent between the dynamical CoTu and CoTuLa models, suggesting that sparse inference in the CoTu model did not suffer from biases in the tuning parameters. Indeed, while there was a statistically significant effect for the PVC neurons (Fig. 7e, inset, P = 1E-13, N = 591, Wilcoxon signed-rank test), its small effect size (Cohen’s d = 0.06) makes its functional impact minimal. No statistically significant effect was observed for the HPC data (Fig. 7e, inset, P = 0.2, N = 467, Wilcoxon signed-rank test, Cohen’s d = 0.004). For the visual cortex data, the result can be intuitively explained because the experimental design ensured that the temporal correlation was minimized in the time-varying stimulus (pseudo-random ordering of orientations). In terms of Eq. 7, we can say that in this experiment, the visual cortex was operating in the *h* ≈ 0 regime, where the tuning biases vanish. For the hippocampus data, on the other hand, tuning biases are not present despite the fact that the animal’s positions are clearly correlated in time (*h >* 0). We next examined the coupling terms (Fig. 7c,f; primary visual cortex and hippocampus, respectively). For the visual cortex data, in the aggregate, the coupling parameters decreased substantially from CoTu to CoTuLa inference procedures (Fig. 7c; inset: P *<* 1E-324, N = 6562, Wilcoxon signed-rank test, Cohen’s d = 0.4). For the hippocampus data, coupling was largely consistent between models (Fig. 7f, inset: P = 8E-9, Cohen’s d = 0.012), and again did not appear to be modulated across the early and late periods. These results indicate that biased inference was pronounced in the visual cortex data, but not the hippocampus data.

**Figure 7:**
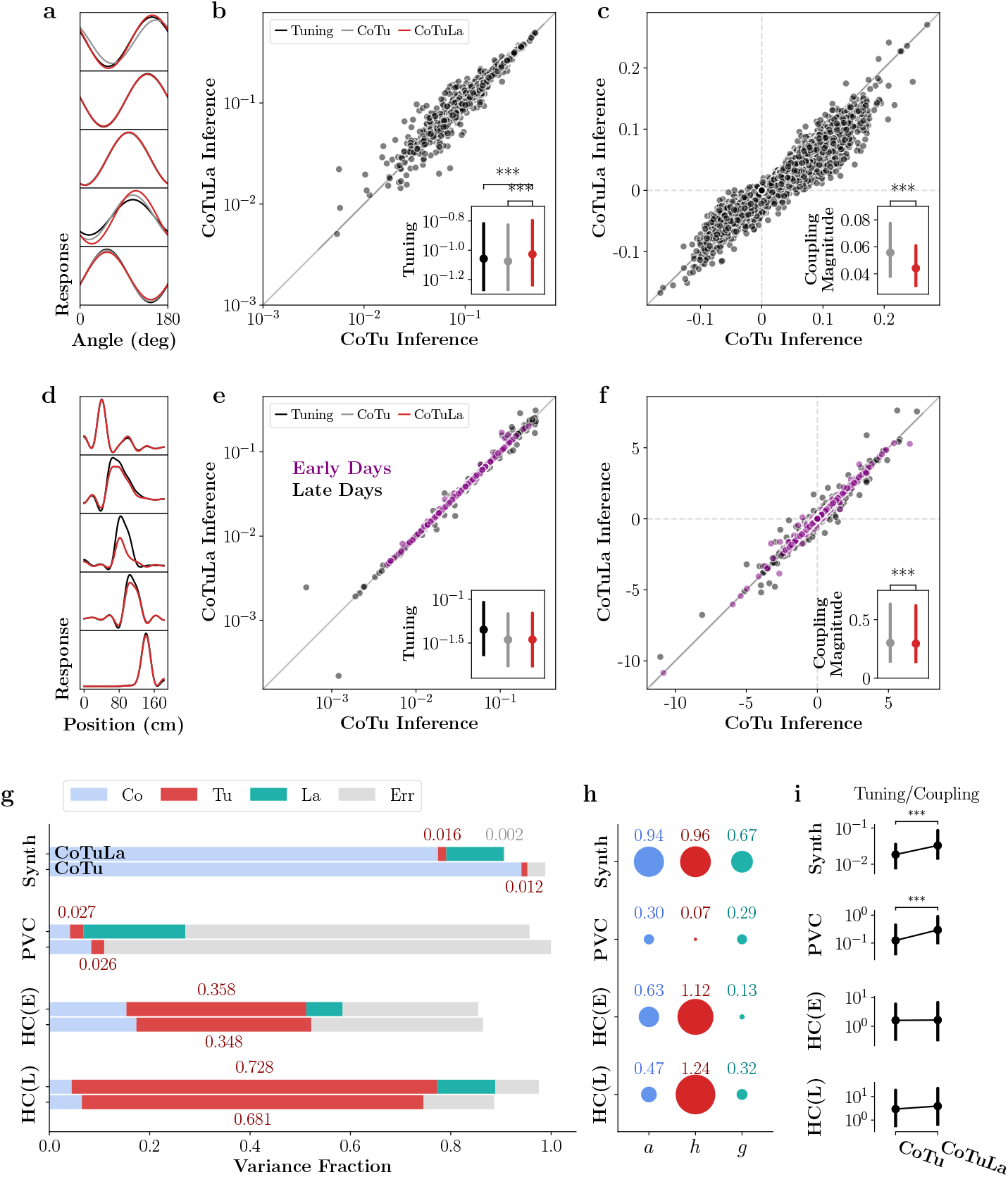
Sparse inference in the dynamical CoTuLa model can mitigate bias in neural data. **a-c**. Data from primary visual cortex. **a**. Example tuning curves from the tuning-only (black), CoTu (grey), and CoTuLa (red) model inference procedures. **b**. Comparison of tuning modulations from CoTu and CoTuLa model inference procedures. Inset: box plots (median and IQR) of the distributions of tuning modulations for the three models. **c**. Comparison of coupling modulations. Scatter plot shows all elements of the coupling matrix. Inset: magnitudes of the non-zero coupling elements (non-zero elements excluded for visual clarity). **d-f**. Similar to a-c, but for data from rat hippocampus. **g-i**. Comparison of fits for synthetic, primary visual cortex and hippocampus data. For the hippocampus data, results from early days (E, days 2-4) and late days (L, days 5-9) are shown separately. **g**. Fraction of variance in the neural responses captured by each term, shown is a stacked bar plot. **h**. Estimated strengths of coupling *a*, tuning correlation *h*, and latent correlation *g*. See text for details. **i**. Tuning-coupling ratios, calculated from the variance fractions, based on the CoTu and CoTuLa model inference procedures. Box plots show the median and the IQR.

To understand the causes of these observations, we examined the other components of the model. We quantified the relative contributions from the coupling, tuning, and latent terms (Fig. 7g) (See Methods Section 5.6 for further details). Note that the variance fractions do not necessarily sum to 1 because the terms in the model are not all orthogonal. We started by revisiting the synthetic data where the stimulus and the noise were temporally correlated by design. The coupling contribution (Fig. 7g, blue) decreased in the CoTuLa model compared to the CoTu model, and the tuning contribution (Fig. 7g, red) increased from CoTu to CoTuLa, although the overall tuning fraction is small compared to the coupling (Fig. 7g). We further explained the differences between fitted parameters from the CoTu and CoTuLa models through the lens of the terms in Eq. 5. In Fig. 7h, we show the magnitudes of autoregression values (i.e., coupling coefficients) (*a*), and strength of temporal correlations in the external variable (*h*) and the latent variable (*g*) (also see Eq. 5). Here, *a* is the eigenvalue of *Â* with largest real part, and *g* is extracted from the optimized CoTuLa fit to the data (see Methods Section 5.4). For *h*, we fitted the stimulus data to a separate VAR_1_ process to obtain the autoregression matrix *Ĥ*, and reported the eigenvalue *h* of *Ĥ* with largest real part. In the synthetic data, this showed that *a, h, g* were all large. This is to be expected from the design of the data generation process, and confirms that the summary quantification of *a, h, g* is capturing salient structure in the data generation process. Finally, in Fig. 7i, we show how the tuning-to-coupling ratio (calculated from the variance fractions) changes from CoTu to CoTuLa models. In this synthetic data set, since tuning is elevated and coupling is decreased from the CoTu to the CoTuLa model (Fig. 7g), the tuning-coupling ratio increases from CoTu to CoTuLa significantly (Fig. 7i; P = 1E-82, N = 750, Wilcoxon signed-rank test; Cohen’s d = 0.4). Thus, we can explain differences in the fitted values of coupling and tuning between the CoTu and CoTuLa models from the terms in Eq. 5.

We next applied this analysis to the real neural data sets. In the visual cortex data, inference in the CoTuLa model resulted in a substantially reduced coupling contribution (Fig. 7g, blue) relative to the CoTu model, suggesting that unlike the tuning parameters (red), coupling parameters were biased when the latent term was not considered. This observation is consistent with the finite *g*, and the small *h*, estimated from this dataset (Fig. 7h). The tuning-coupling ratio increased significantly from CoTu to CoTuLa, due to the decrease in the coupling contributions (Fig. 7i; P = 5E-95, N = 591, Wilcoxon signed-rank test; Cohen’s d = 0.4). Our results further indicate that there was a substantial amount of unobserved, temporally correlated variability in the primary visual cortex recordings in this dataset (Fig. 7g, green). We did not observe any notable difference between the two monkeys in this data set. These results indicate that even with temporally uncorrelated stimuli, failure to consider the effect of correlated unobserved variability may lead to misleading conclusions about the relative importance of tuning and coupling terms in the neural system, especially if there are systematic differences in the strength or temporal correlation of unobserved variability, as might occur during learning.

In the hippocampus data, both coupling and tuning contributions were largely unaffected by the inclusion of the latent term (Fig. 7g, blue and red), suggesting that there was neither coupling nor tuning biases in the CoTu fit (consistent with Fig. 7e-f). The relatively small value of *g*, especially in the early days (days 2-4) of the experiment, is consistent with this observation (Fig. 7h). The tuning-coupling ratio changed little in both cases (Fig. 7i). Interestingly, compared to the early days, in the later days (days 5-9) of the experiment, the overall variance fraction was more heavily influenced by tuning compared to coupling (Fig. 7g, red and blue). Furthermore, there was a substantial increase in the contribution of variance attributed to the unobserved latent variable (Fig. 7g, green), and the temporal correlation of the variability *g* was larger (Fig. 7h). This may be related to the fact that the rat was more familiar with the task environment in the later trials of the task. To examine this in more detail, in Fig. 8a, we visualized the variance fraction for the different components (different colors) of both models (CoTu: dashed lines; CoTuLa: solid lines) as a function of the number of days the rat has been in the experiment. We observed substantial modulations of the tuning (red) and coupling (purple) parameters in both models as a function of exposure to the experiment. In particular, tuning and coupling were nearly on par early on (e.g., day 2,3), but tuning quickly become the dominant factor compared to coupling (days 4-5). Coupling and tuning largely stabilized after day 5. The latent variable in the CoTuLa model (teal) appeared to be anti-correlated with the tuning parameter in the later days. Statistical tests of comparisons between early (purple) and late (black) days of the variance fractions for coupling, tuning, and latent variables are provided Fig. 8b. Together, these results clearly reveal how the severity of biases depends on the characteristics of the experimental design with which the data set was collected, and on the details of the system (i.e., the strength of coupling, temporal correlation of unobserved variability) that generated the data.

**Figure 8:**
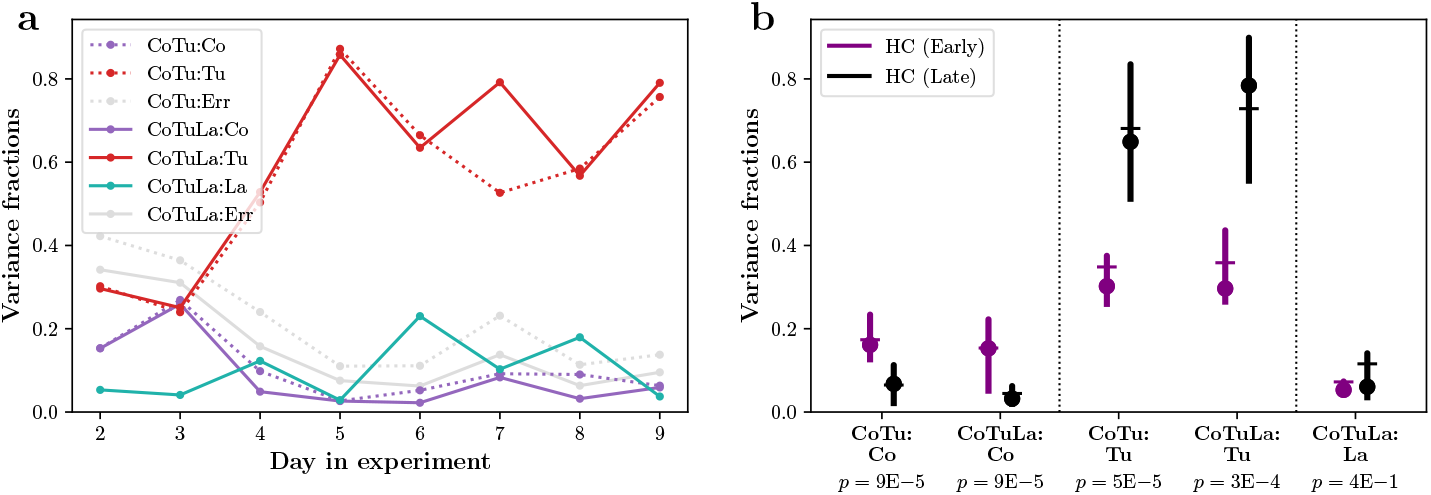
Modulation of tuning, coupling and latent parameters across early and late phases in hippocampal experiment. **a**. Day-by-day view of the variance fractions, showing large changes over the first few days of the experiment. **b**. Comparison of variance fractions between the early phase (days 2-4 since the animal was introduced to the experiment) and the late phase (days 5-9). Box plots show median (dot), mean (horizontal line) and the IQR. P values are from Wilcoxon rank-sum tests comparing distributions between early and late phases.

### 3.7 Sparse model support resolves structural non-identifiabilities in the static CoTuLa model

In the results above, we consistently observed that complete removal of bias requires not only specification of the static/dynamic CoTuLa model, but more interestingly, on accurate inference of model support (i.e., variable selection) before parameter estimation in those models. That is, even when the structure of the model is correct, inference in that model is biased without first constraining the support of the model. Why is this the case? To answer this question, we return to consideration of the static CoTuLa model. The static CoTuLa model, as defined in its linear-Gaussian form, suffers from structural non-identifiability [25]. Structural non-identifiability in a probabilistic model exists when an infinite number of transformations can be applied to any parameter configuration that results in a new parameter configuration that has equal likelihood. This is visualized in the example 2D loss surface of (Fig.9a), where the red lines indicate examples of parameter space (i.e., parameter 1 and parameter 2) that have the same loglikelihood values. The existence of an infinite number of parameter values with the same value of loss impairs the interpretability of the fitted model parameters, since for any inferred set of parameters on a likelihood surface (e.g., obtained via EM optimization), one can always find a family of parameter estimates that have the same log-likelihood (Fig. 9a). Structural non-identifiability has been studied in systems biology, and in some neuroscience models [26, 27]. However, the consequences of structural non-identifiability on fitted parameters are poorly understood, and approaches to render a non-identifiable model identifiable are nascent.

**Figure 9:**
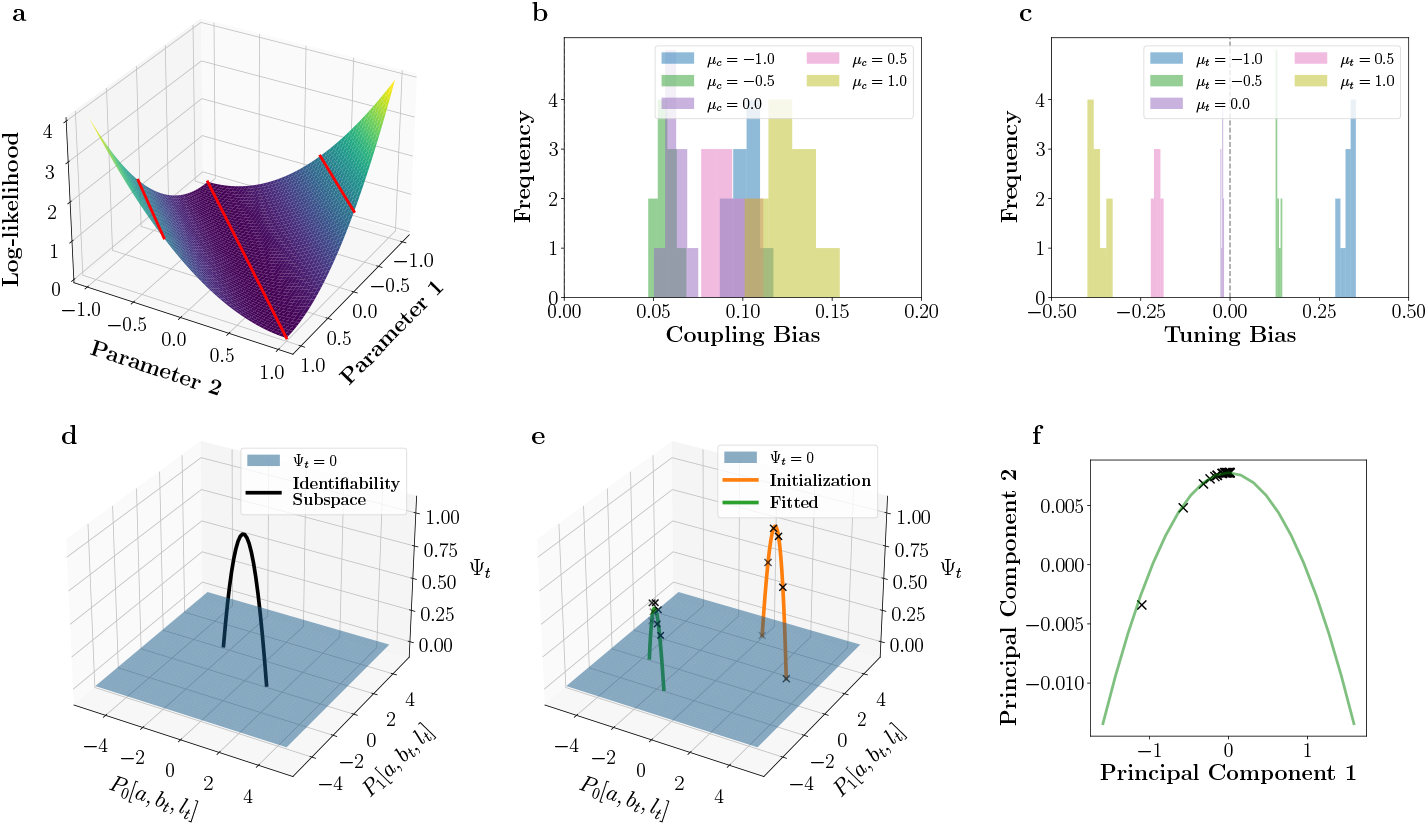
Structural non-identifiabilities within the static CoTuLa model contributes to consistent biased estimation. **a**. A toy loss surface (*z*-axis) in two parameters (*x*and *y*-axes). Red lines denote example identifiability subspaces for this loss surface. **b-c**. Biases calculated for coupling and tuning parameters obtained from CoTuLa inference on data generated from a sparse, identifiable CoTuLa model. CoTuLa inference was conducted without specification of a model support, and thus had structural non-identifiability. Biases are shown across 30 different initializations. Different colored histograms correspond to different means of the underlying parameter distributions. **d**. The identifiability family for the static CoTuLa model behaves like a truncated parabola when the latent dimension *K* = 1. Black line denotes the identifiability family plotted as a function of principal axes *P*_0_ and *P*_1_, which depend on *a, b*_*i*_, and *l*_*i*_. The black curve denotes parameter configurations for which the loglikelihood is unchanged. The blue surface denotes the point of truncation, where the private variance Ψ_*i*_ would become negative. **e**. The experiment set-up for examining structural non-identifiabilities. Orange curve denotes an identifiability family at initialization. Many models are initialized at different points along the identifiability family (denoted by ×). Parameter inference is performed using EM until a fitted identifiability family is reached (green curve). The fitted solutions (denoted by × on the green curve) cluster near the top of the identifiable class, where private variance is maximized. **f**. The fitted solutions (× marks) obtained empirically from the experiment described in **e**., but visualized with two principal components.

Intuitively, structural non-identifiability may have been expected since the number of unobserved neurons in many neuroscience experiments is much larger than the number of recorded neurons. Without further specification, the general problem of inferring tuning and coupling parameters with many unobserved neurons that have a strong functional influence on the observed neurons is likely intractable [8, 28]. However, our results indicate that the actual overall effect of unobserved neurons is modest in real neural data. Furthermore, we have decades of consistent experimental observations about neural data that may constrain the general problem. Indeed, despite these theoretical challenges, we identify two neurally plausible conditions under which the static CoTuLa model is identifiable. Namely, we next show that if the data generating system has sparse enough tuning and coupling parameters and the latent influence has low enough dimensionality, then the static CoTuLa model is identifiable. Sparse coupling and tuning are commonly used as a prior to improve functional coupling and tuning estimates [19, 29, 30]. Interarea neural communication and spontaneous activity due to task-irrelevant behavior have also been shown to be relatively low-dimensional compared to the ambient neural dimensionality [31, 32]. Thus, given the known structure of neural data, the generally intractable problem can become tractable.

We first analytically identified the nature of the structural non-identifiability in the static CoTuLa model. We specify an identifiability transform, a transformation that can be applied to any set of parameters in the CoTuLa model to obtain another set of parameters with equal log-likelihood of data, via a tuneable parameter. We call the set of parameter configurations reachable via an identifiability transform an identifiability family (Fig. 9a, individual red lines correspond to different identifiability families). The identifiability family in the CoTuLa model is linear in the target tuning, coupling, and latent factor parameters, and quadratic in the target private variance. It is specified by the *K*-dimensional vector ***δ***, with transform 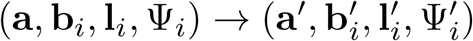 given by

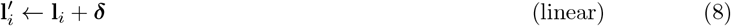

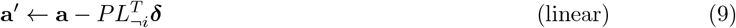

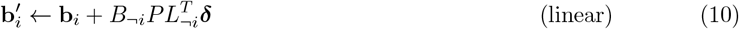

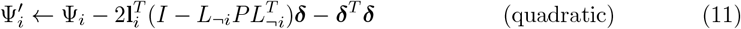

where 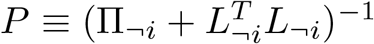, i.e., the neural variability precision matrix. Here, (from 2 and 3), **a** and **b** are the coupling and tuning parameters, respectively, and **l** and **Ψ** are the shared and private components of the unobserved variability, respectively.

We determined the specific conditions under which sparse support in the coupling and tuning parameters could remove the structural non-identifiability present in the CoTuLa model. We show the following: given a procedure for sparse estimation of **a** and **b**, there is a required level of sparsity in **a** and **b**_*i*_ (and an additional condition on the rank of a matrix) such that any identifiability transform modifies the support of **a** and **b**_*i*_ and so support-preserving estimation is identifiable. We provide the specific conditions in Theorem 1, for which we provide a proof in Methods Section 5.7.2. The basic intuition is that sparse feature selection constrains the model so as to remove non-identifiability.

**Theorem 1**. *Consider an identifiability transformation of the tuning and coupling parameters:*

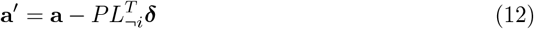

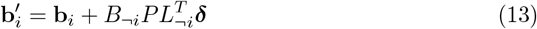

*where there are N coupling parameters in* **a**, *M tuning parameters in* **b**_*i*_, *and K latent factors*. Π_¬*i*_, *L*_¬*i*_, *and B*_¬*i*_ *are fixed. Let k*_*C*_ *be the sparsity of* **a** *so that Nk*_*C*_ *elements of* **a** *are exactly zero. Similarly, let k*_*T*_ *be the sparsity of* **b**_*i*_ *so that Mk*_*T*_ *elements of* **b**_*i*_ *are exactly zero. Let* 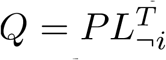 *and R* = *B*_¬*i*_*Q and let Q*_*sub*_ *and R*_*sub*_ *be their respective matrices only including the rows that are not in the supports of* **a** *and* **b**_*i*_, *that is, only including the rows corresponding to values of 0 in* **a** *and* **b**_*i*_. *Recall that P is the neural variability precision matrix, so Q*_*sub*_ *is a projection of the shared variability onto the precision matrix, and R*_*sub*_ *further projects this into the non-target tuning*.

*If K Nk* + *Mk and* 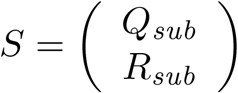 *is full-rank, then the only* ***δ*** *in the identifiability subspace which preserves the model support of* **a** *and* **b**_*i*_ *is* ***δ*** = 0.

This theorem shows that parameter estimation can be identifiable given sparse enough parameters, and that *S*, a linear transform of the unobserved precision matrix, is full rank. In practice, sparse model fitting can often be split into two separate steps: support recovery and non-zero parameter estimation [18]. Here, we have shown that estimating the non-zero parameters is identifiable, although accurate support recovery in correlated, noisy datasets is still an area of active research [33, 34]. None-the-less, we have turned a problem that was completely intractable into one that is tractable, though admittedly hard.

### 3.8 Structural non-identifiability in the static CoTuLa model contributes to consistent biased estimation

Structural non-identifiability impacts parameter estimates and model interpretability if not resolved prior to parameter inference. For example, suppose we fit a static CoTuLa model to data generated from an identifiable model with sparse support, in accordance with Theorem 1. Since the ground truth model parameters satisfy Theorem 1, there is a unique parameter configuration that generated the data. Suppose, however, we fit a CoTuLa model without enforcing the conditions of Theorem 1 (e.g., not requiring sufficient sparsity during parameter optimization). In this scenario, where the *inferred* model is structurally non-identifiable, how are the resulting parameter estimates impacted? As there are an infinite number of parameter values with the same log-likelihood of observing the data (i.e., the identifiability family), one common intuition is that estimated parameter values will be highly variable across multiple runs of the inference algorithms. In the context of EM, this could translate into extreme sensitivity to the initial conditions. However, the analytic results for the dynamic CoTuLa model revealed biases that depended on the data generation process, and the numerical results of both the static and dynamic CoTuLa models showed biased estimates but low variability. Thus, in contrast to common intuition, it appears that, in these cases, non-identifiability resulted in consistent estimation of the wrong answer, not highly variable estimates. To gain further insights into how non-identifiability impacts bias and variance of estimated parameters, we assessed the impacts of fitting structurally non-identifiable models to data generated from identifiable ground truth parameters.

We conducted a simple experiment in which we fit a structurally non-identifiable CoTuLa model to data generated from a sparse, structurally identifiable static CoTuLa model. To simplify exhibited bias, the data generation model had positive tuning parameters, as is the case for neural data. Given that the fitted model was structurally non-identifiable, there existed an entire identifiability family on parameter estimates that have equal values of the EM objective function. We calculated coupling and tuning parameter biases obtained from dense parameter inference (i.e., without first imposing model support) across 30 random initializations of EM. The distributions of these biases across initializations are depicted in Figure 9b-c. We also varied the mean of the distribution from which the model parameters were drawn (different colored histograms). We observed that coupling parameters are overestimated (Fig. 9c, strictly positive bias across parameter distributions with different means), while the magnitude of tuning parameters are underestimated (Fig. 9c; bias is opposite in sign from the mean of the parameter distribution, indicating a reduction in magnitude). These biases are qualitatively similar to Fig. 3. Interestingly, we observe low variance across initializations. Despite the fact that inference was conducted on a structurally nonidentifiable model–where a wide range of parameter values are equally valid under the log-likelihood–parameter inference consistently resulted in similar estimates across initializations. The resulting estimates are a high-bias, low-variance estimate of the ground truth parameters.

The low variance observed in Figure 9b-c is surprising given that we might expect a wide range of parameter estimates, spanning an identifiability family. Since the identifiability family is linear in all parameters except the target variance (Equations 8-11), the high-dimensional identifiability subspace can be visualized as a truncated parabola (Fig. 9d: black curve). The parabola is truncated when the private variance (Ψ) reaches zero at either end (Fig. 9d: blue surface). Thus, the low variance we observe on the fitted parameters implies that these solutions are clustering on some location of the truncated parabola.

It may be possible that the random initializations during inference are small in the geometric context of the identifiability family. Thus, we repeated the experiment, but with a different set of initializations: those that span an initial identifiability family. These can be thought of as the initializations with the largest amount of variance, with respect to the identifiability family. We chose a random initialization and calculated the identifiability family around it (Fig. 9e: orange curve). We then chose 30 equally spaced parameter configurations spanning the identifiability family as initializations for parameter inference (Fig. 9e: x’s on the orange curve). We conducted parameter inference from each of these points in the non-identifiable model. We hypothesized that, if structural non-identifiability did not impact parameter bias, then the inferred parameters would similarly lie equally spaced on an identifiability family. In contrast, if structural non-identifiability imparts bias, then the inferred parameters would be consistently clustered around a particular region of the identifiability family. To visualize the results, we performed principal components analysis on the fitted parameter estimates across fits, plotting the space in two dimensions (Fig. 9f). The fitted solutions clustered around the apex of the parabola, implying that, despite significantly different initializations, optimization resulted in a set of parameter estimates with low-variance. Interestingly, this solution was that which maximized the target private variance (i.e., the apex of the parabola, which corresponds to private variance). These results demonstrate that failure to resolve structural non-identifiability may produce biases during parameter inference, despite the use of the correct underlying model (in this case, the model actually that generated the data). Thus, accurate selection in a structural accurate model is crucial in achieving low-bias parameter estimates.

## 4 Discussion

We have identified an underappreciated bias in parameter estimates arising from structural non-identifiability in models of functional coupling amongst neurons and the tuning to external variables. Similar bias was present in both static and dynamic models, and persisted with the inclusion of latent sources of variability in the model. We demonstrated that bias was mitigated by specifying accurate model support in both the static and dynamic cases; in the static case, we proved that accurate model support removes model non-identifiability. Application of models and inference algorithms to diverse neural data sets revealed heterogeneity in the influence of coupling to other neurons relative to tuning to external stimuli. Below, we discuss these results from a (neuro)-biological and statistical perspective.

### 4.1 Interactions, externals, and unobservables in (neuro-)biology

Many biological processes engage the interaction of multiple components that are themselves affected by external factors. For example, the integrity and function of the rhizosphere (the top layer of soil surrounding plant roots), is maintained through the interaction of many microbes, which themselves are impacted by soil conditions (e.g., nitrogen and water levels), as well as root exudates. Likewise, in the brain, it is the interaction of many neurons that produce sensation, cognition, and behavior, and neuronal activities are often modulated by sensory inputs (e.g., sounds, sights, etc.). In all but the most controlled experiments, it is not possible to simultaneously measure all the factors that influence the activities of the, e.g., neurons we are trying to understand. Determining if and how different biological factors influence observed cellular activities is central to our understanding of complex biological systems. In many cases, this understanding must be extracted from statistical modeling of the data.

In the specific context of statistical models of neural activity, we determined if, and to what extent, estimates of parameters corresponding to interactions (i.e., functional coupling) and external variables (i.e., sensory tuning) were affected by biased inference. To account for the effect of unobserved neurons in recorded data, we extended the CoTu model of neural activity with inclusion of latent variables (CoTuLa). We also introduced a two-stage algorithm to first infer the non-zero parameters, and then estimate the values of those parameters. Both the inclusion of a latent variable and a two-stage inference algorithm were found to be important to mitigate bias in synthetic data, with analogous changes in parameter estimates observed in diverse real neural data.

We examined four neurophysiology data sets, one from auditory cortex (rat), two from visual cortex (monkey), and one from the hippocampus (rat). In all three of the cortical data sets, we observed that the CoTuLa model with two-stage inference significantly and substantially increased the inferred contribution of tuning relative to coupling compared to the CoTu model. These observations, combined with our numerical and analytic investigations, strongly indicate that standard inference in the CoTu model was substantially biased. Nonetheless, these results corroborate previous findings (using the CoTu model and basic inference) that coupling to other neurons is the dominant source of variability in cortical neuronal activity [8]. Interestingly, for the static models, we found that coupling was a more dominant factor for the single unit recordings than for the ECoG recordings. This may seem surprising, as ECoG records electrical fields from the cortical surface and is thought to have low spatial resolution. In contrast, single-unit recordings measure the action potentials of putative single neurons, which therefore have single-cell resolution. However, recent results indicate that the high-gamma component of the ECoG signal analyzed here is localized to single cortical columns and reflects population firing rates of supragranular neurons [35]. Further, the *μ*ECoG grids utilized in these experiments had 200 *μ*m spacing (3.2×1.6mm total area), and span multiple auditory cortical regions of the rat brain [35]. In contrast, recordings from the visual cortex were performed in monkeys with arrays with 400 *μ*m spacing (3.6×3.6 mm total area), but were likely contained to the primary visual cortex [23]. Anatomical neuronal connectivity is typically denser and stronger within an area than across areas [36]. To the degree that estimates of functional coupling are reflective of anatomical connectivity, the observed differences in coupling contributions may reflect the differences in spatial coverage of functional areas by the recording devices (other differences in the experimental settings notwithstanding).

The results obtained from the rat hippocampus data are qualitatively different than the results described above for data recorded from primary sensory cortices. In the sensory cortex data, CoTu-biased inference enhanced the importance of coupling relative to tuning, and coupling was the dominant source of variance. In contrast, in the hippocampus data, we found minimal evidence of CoTu-biased inference, and tuning was the dominant source of variance compared to coupling (in contrast to [8]). These heterogeneous effects across the data sets may be explained by differences in the structure of latent variables (as well as the nature of the neural code); for example, correlated variability was largest in magnitude and most uniform in sign for the ECoG data, smaller in magnitude and more heterogeneous in sign for the PVC data, and smallest and most heterogeneous in the hippocampus (c.f., Fig.1). This ordering of correlated variability across data sets mirrors the ordering of observed CoTu-bias. As attention and learning can modulate correlated variability [37, 38], it would be important to properly account for the issues revealed here when interpreting the effects of learning and attention on tuning and coupling. We found pronounced effects of learning on the relative magnitudes of coupling and tuning (but not latent) contributions to hippocampus neuronal activity. In particular, the dominance of tuning increased (to asymptote) with learning, with an accompanying decline in the contribution from coupling. A previous examination looked at the overall levels of coordinated activity as a function of novelty and behavioral state in [39]. It was found that correlated firing was much more prevalent early in learning while place tuning could explain more of the data later in the task. Our results agree with that previous observation, and further refine the source of correlated firing to be couplings amongst neurons, not unobserved latent influences.

In all data sets, the recorded neuronal activity is likely modulated by neurons that are not being monitored. The incorporation of latent variables acts as a way to account for omitted neurons in the model. The inclusion of other omitted variables that are not accounted for by the latent variable, such as additional tuning parameters (e.g., the spatial frequency of the gratings), may also be important in order to extract unbiased and interpretable parameter estimates from the model [12]. In this work, the latent variables that generated correlated variability were independent of any task, stimulus, or neural state variables. However, correlated variability has been shown to have stimulus and task dependence [22, 40]; that is, the covariance structure of neural response around any fixed stimulus depends on that stimulus. To the extent that correlated variability can be modulated through task-structure (attention) or measured (spontaneous behavior), CoTuLa models where the latent variables are conditioned on these factors could be developed and estimated from data [32, 38, 41, 42].

Our analysis of the dynamic model indicates that parameter bias should become more pronounced as the temporal extent and magnitude of correlations in behaviors and/or sensory stimuli increase. There is a well-motivated movement in systems neuroscience to engage animals in more complex, ethologically relevant tasks [43], which may have longer temporal correlations than classical experimental paradigms. Thus, the potential for substantial parameter bias may increase in the future. It has been argued that in the context of substantial contributions from unobserved neurons, inferring functional coupling that accurately reflects anatomical connectivity is intractable [8, 28]. The intuition being that functional coupling is effectively a correlational measure based on firing rates, and if the correlations in firing rates are dominated by unobserved neurons, inference of functional coupling is doomed. However, we observed minor to modest-sized contributions of latent terms in our models. Thus, as increasingly large numbers of neurons are being recorded simultaneously from across the brain, many of the known issues with functional coupling estimates (in particular, sparse sampling of the neurons) are likely to subside. Therefore, robust methods for estimating the interactions amongst recorded neurons are well poised to have a sustained impact in the future. Such extracted networks could, e.g., be used to analyze network controllability [44], etc., to guide closed-loop perturbations of neural activity.

### 4.2 All models are wrong, but some are useful

“All models are wrong, but some are useful.” This adage from George Box serves as both warning and inspiration for statistical modeling, especially in the biological sciences. One way a model can be useful is the degree to which it predicts the results of new experiments. This notion of model use is, in part, the motivation for using metrics such as cross-validated predictive accuracy for model evaluation. However, in basic biology, it is rarely the case that a model trained on one data set is used to predict the results of other, new experiments quantitatively. Another way a model can be useful is the degree to which it provides insight into the processes that generated the observed data to which the model is fit. A quantitative treatment of this measure of usefulness necessarily entails analysis of the parameters of the model. Examples of this use of models abound across biology: fitting parametric tuning curves to single-neuron activity in response to a stimulus [45]; fitting Lotke-Voltera models to microbial data to understand their dynamic interactions [46] or fitting models of tuning and coupling to population neural data to ascertain the relative importance of those influences on neural activity [12]. Indeed, we argue that insight is often the most important use of statistical modeling for exploratory, data-driven discovery in biology.

The use of statistical models to provide insight into biological processes hinges on the quality of the inferred parameters. However, there are many ways in which the parameters of a model can be misleading. It is useful to briefly summarize such failure modes, as well as modern approaches to address them, as our results reveal previously unappreciated interactions between these failure modes, and many modern solutions may be unknown to the broader computational biology community. We ground our discussion in the context of linear regression, as this is where these concepts are most intuitively understood.

The most well-known form of parameter error is the bias-variance trade-off, which formalizes the difficulty in minimizing both the bias (inverse of accuracy) and variance (inverse of precision) contributions to parameter error during inference [47]. Variance can presumably be identified through examination of parameter estimates resulting from various re-sampling approaches (e.g., bootstrap resamples, resampling of initial conditions of an inference algorithm, etc.), and potentially mitigated through the aggregation of such results [48]. High variance can result from, e.g., numerical issues associated with ill-conditioned inverse covariance matrices. We assert that parameter bias is a much more detrimental and insidious form of parameter error than variance, as it is typically more difficult to asses via standard numerical approaches (e.g., resampling). That is, if the parameter estimates are consistent and one does not evaluate the procedure relative to a known ground truth, how would one ever know if the estimates are consistently wrong in small samples? As we have shown, the sources of parameter bias can be nuanced and challenging to resolve.

Another form of parameter error that receives somewhat less attention (though it played a central role in this work) is the inferential accuracy of model support (i.e., parameter or model selection). As with parameter estimation, there are two forms of error here, false-positives and false-negatives, and the goal of an inference algorithm is to achieve low error rates in both. Roughly speaking, both the false-positive and false-negative rates will depend on the smallest signal-to-noise in the parameters and the amount of multi-collinearity amongst the parameters [18]. Inference of model support, i.e., imposing sparsity, is commonly achieved through the inclusion of structured regularizers to constrain the optimization. While _1_-regularization (as targeted by, e.g., Lasso [17]) is the most commonly deployed approach, it suffers from biased parameter estimates and low selection accuracy [18, 33, 49, 50]. Regularizers that (asymptotically) achieve oracle support recovery and have low bias have been introduced, such as SCAD and MCP [49, 50]. However, because these algorithms formulate non-convex optimization problems, they can suffer from a relatively high variance of parameter estimates [51]. Recently, we have introduced Union of Intersections (UoI), a two-stage inference framework which achieves nearly oracle support recovery with nearly unbiased and low-variance estimates [18]. In practice, the difference in support recovery between SCAD, MCP, and UoI is negligible [33].

As discussed above, the typical way in which we think about the interaction of these various forms of parameter error (bias, variance, false-positive, false-negatives) during inference comes in through the inclusion of structured regularizers to constrain the optimization problem [52]. However, if and how these types of errors interact in the context of the structure of a model is less well understood. As we and others have shown, the interpretation of a model is greatly hindered by the structural non-identifiability of the model itself. Put bluntly, if there is an infinite family of model parameters that are equally good at predicting observed data, how can we trust the parameters we get? The first-order intuition is that structural non-identifiability would be evinced by high variance estimates of model parameters; the final parameter estimates (i.e., where in the identifiability family optimization ends) could depend strongly on the initial conditions [47]. Despite the intuitive appeal of this line of thinking, our results indicate that, in the context of the (static and dynamic) CoTuLa model studied here, inferred parameters exhibit high bias and low variance. That is, parameter optimization terminates at the same location in the identifiability family when starting from random initial conditions, and that final location is biased relative to the parameters that generated the observed data.

In the static CoTuLa model, we showed that estimates from *ℓ*_1_-regularized EM are also biased, suggesting that arriving at unbiased estimates requires resolving structural non-identifiability before inferring model parameter estimates. During optimization of an *ℓ*_1_-constrained problem (or any other structured regularizer, for that matter), the initial parameter space is fully dense, as it is only through the traversal of an optimization trajectory that parameter sparsity is introduced. This implies that the initial optimization steps will be dictated by the gradient derived from the non-identifiable model. These considerations together with our results suggest that it is the initial steps that take optimization along a trajectory that terminates in biased parameter estimates. Once along this trajectory, a gradual introduction of sparsity does not appear to bring optimization to an unbiased solution. We note that the requirement of true sparsity (i.e., parameters not in the support are set exactly to 0) to resolve structural nonidentifiability before parameter estimation indicates that Bayesian approaches, which typically require a *post-hoc* procedure to set parameters exactly to 0 [53], will not resolve these issues.

By identifying that accurate model support resolves model structural non-identifiability and can result in accurate parameter estimates, we have taken a mathematically degenerate problem (inference in non-identifiable models) and turned it into a problem that is tractable and for which concrete progress can be made. Importantly, however, we emphasize that our understanding of the relationship between parameter bias and structural non-identifiability, while empirically quantitative, is not theoretically analytical. If the observations made here can be proven theoretically, it would argue that two-stage procedures that specify model support and then perform estimates just for the non-zero parameters, such as those deployed here, are strictly required for accurate parameter estimation in this class of models. We believe that arriving at a deep theoretical understanding of this issue is an interesting statistical problem in and of itself, and one that requires novel theoretical work that is outside the scope of this manuscript.

## 5 Methods

### 5.1 Neural recordings and data analysis

#### 5.1.1 Recordings from auditory cortex

Auditory cortex (AC) data was comprised of cortical surface electrical potentials (CSEPs) recorded from rat auditory cortex with a custom fabricated micro-electrocorticography (*μ*ECoG) array. The *μ*ECoG array consisted of an 8 × 16 grid of 40 *μ*m diameter electrodes. Anesthetized rats were presented with 50 ms tone pips of varying amplitude (8 different levels of attenuation, from 0 dB to −70 db) and frequency (30 frequencies equally spaced on a log-scale from 500 Hz to 32 kHz). Each frequency-amplitude combination was presented 20 times, for a total of 4200 samples. The response for each trial was calculated as the *z*-scored, to baseline, high-*γ* band analytic amplitude of the CSEP, calculated using a constant-Q wavelet transform. Of the 128 electrodes, we used 125, excluding 3 due to faulty channels. Data was recorded by Dougherty & Bouchard (DB), and can be found in Collaborative Research in Computational Neuroscience (CRCNS) data sharing website [54] as [55]. Further details on the surgical, experimental, and preprocessing steps can be found in [35].

#### 5.1.2 Recordings from primary visual cortex

We analyzed primary visual cortex (V1) datasets, comprised of spike-sorted units simultaneously recorded in anesthetized macaque monkeys. Recordings were obtained with a 10 × 10 grid of silicon microelectrodes spaced 400 *μ*m apart and covering an area of 12.96 mm^2^. In the static experiments, three monkeys were presented with grayscale sinusoidal drifting gratings, each for 1.28 s. Twelve unique drifting angles (spanning 0° to 330°) were each presented 200 times, for a total of 2400 trials per monkey. Spike counts were obtained in a 400 ms bin after stimulus onset. We obtained [106, 88, 112] units from each monkey. In the dynamic (gratings movie) experiments, two monkeys were presented with similar drifting gratings, but with changing orientations every 300 ms. Spike counts were binned at 300 ms intervals that match stimulus changes. We obtained [74, 123] units from each monkey. The data was obtained from the Collaborative Research in Computational Neuroscience (CRCNS) data-sharing website [54] and was recorded by Kohn and Smith (KS) [23]. Further details on the surgical, experimental, and preprocessing steps can be found in [56] and [57].

#### 5.1.3 Recordings from hippocampus

We analyzed rat hippocampus (CA1 and CA3) datasets, with spike-sorted unit activities from rats while the animals perform an alternate choice task in W-shaped tracks. We analyzed data from a single animal named “Bond”. Of the 24 available data epochs (over 8 days) from this animal, we used 16 epochs (2 epochs from each day) where the animal was put in the same track. The four early days (8 epochs) and the four late days (8 epochs) were analyzed separately. We chose cells with firing rates ≥ 0.1 Hz, obtaining [413, 359] units from early and late subsets. Firing rates were obtained by binning at 200 ms. The data was obtained from the CRCNS data sharing website [54] and was recorded by Karlsson, Carr and Frank [24]. Further details of the experiments can be found in the original papers [58, 59].

#### 5.1.4 Modeling tuning with basis functions

We used basis functions to model the influence of tuning in the neural data. We use a set of *M* basis functions *g*_*i*_(*s*), which form the *M* -dimensional stimulus representation **x**. The contribution to the activity of a specific neuron provided by the stimulus is encoded by tuning parameters **b**, where

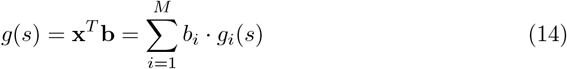

For the primary visual cortex data, we use cosine basis function (using *M* = 2). For the auditory cortex data, we used Gaussian basis functions. Specifically, we chose *M* = 8 basis functions tiling the log-frequency plane with a standard deviation of *σ*^2^ = 0.64 octaves. For the hippocampal data, we used *M* = 10 Gaussian basis functions tiling the linearized positions of each trajectory, with a standard deviation of 10cm.

### 5.2 Testing on synthetically generated data

#### 5.2.1 Synthetic data generation for the static model

We chose *N* = *M* = 10 coupling and tuning parameters, setting half of them equal to zero (e.g., a sparsity of *k*_*T*_ = *k*_*C*_ = 0.5), and a noise correlation of *ρ*_*C*_ = 0.25 with *K* = 1 latent factor. Given the tuning and coupling parameter means (the hyperparameters varied in the experiment), we drew both tuning parameters and coupling parameters from Gaussian distributions, with variance *σ*^2^ = 0.5.

#### 5.2.2 Synthetic data generation for dynamical model

The observed data *Y* was generated according to **y**_*t*_ = *A***y**_*t*−1_ + *B***x**_*t*_ + ***ϵ***_*t*_, where ***ϵ***_*t*_ 𝒩(**0**, Σ). The *N* × *N* coupling matrix *A*, was drawn element-wise from a log-normal distribution (mean 0 and standard deviation 1 for the underlying normal distribution), then sparsified by randomly dropping out half of the elements to zero (density 0.5). If the largest eigenvalue *λ*_max_ of *A* was larger than 0.95, the matrix was scaled by a constant factor 0.95*/λ*_max_ to ensure stability. The *N* × *M* tuning matrix *B* consisted of a set of Gaussian bumps, with a Gaussian bump in each row, with fixed peak amplitudes (1.0) and widths (1.2 columns). The peak position was varied in a space-tiling fashion, such that the peaks form a band from the top left of the matrix to the bottom right. The *N* × *K* latent coupling matrix *L* had only one 1 in each row, and otherwise 0. In the case shown in the paper, we used *K* = 1, so *L* was simply a *N* × 1 matrix of all 1’s. The *N* × *N* noise covariance matrix 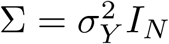 was a uniform diagonal matrix with scale *σ*_*Y*_ = 0.1. The external stimulus *X* was generated from a separate VAR_1_ process with the noise covariance matrix 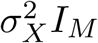, *σ*_*X*_ = 0.1. The coupling matrix was either zero (uncorrelated case) or uniform diagonal *hI*_*M*_, *h* = 0.9 (temporally correlated). The unobserved data *Z* was also generated from a separate VAR_1_ process, with the noise covariance matrix 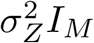, *σ*_*Z*_ = 1.0. The coupling matrix was either zero (uncorrelated case) or uniform diagonal *gI*_*K*_, *g* = 0.9 (temporally correlated). We used *N* = *M* = 15 and *K* = 1. In each realization, we generated 10 time-series (“trials”) with *T* = 100 samples in each trial.

### 5.3 Parametric Inference in the static CoTuLa Model

Since the static CoTuLa model is a latent variable model, we can perform parametric inference via the expectation-maximization algorithm. At the same time, the linear-Gaussian instantiation of the model lends itself well to analytic derivations of the joint and marginal distributions. Here, we calculate these distributions as pre-requisites for deriving the EM-update rules for optimization.

#### 5.3.1 Joint distribution of the neural activity and latent state

A full likelihood expression of the static CoTuLa model incorporates the parameters *L* = [**l**_*i*_, *L*_¬*i*_] and **Ψ** = [Ψ_*i*_, **Ψ**_¬*i*_] that defines the shared and private variability. Recall that the neural activities are defined as

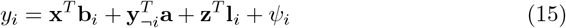

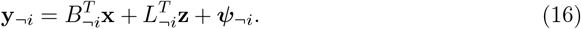

The joint distribution of the data, including the latent variables, can be written as

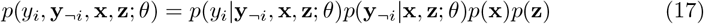

where *θ* specifies the parameter set. In the linear-Gaussian setting, each of these densities takes on the form

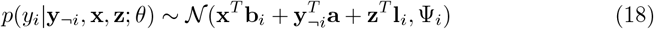

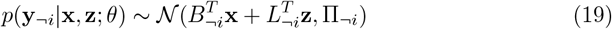

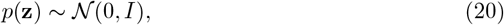

where Π_¬*i*_ := diag(**Ψ**_¬*i*_). By the Gaussianity of the above distributions, we can write the joint distribution of *y*_*i*_, **y**_¬*i*_, and **z** (conditioned on **x**) as a multivariate Gaussian distribution. Specifically, we have

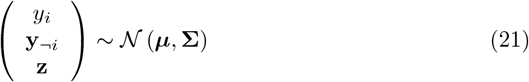

where

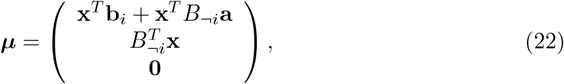

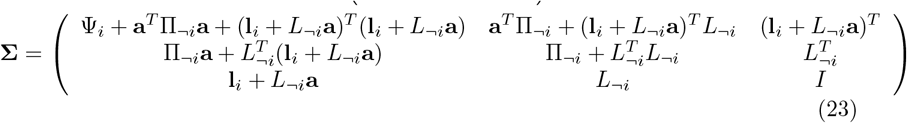

and an associated precision matrix with the analytic form given by

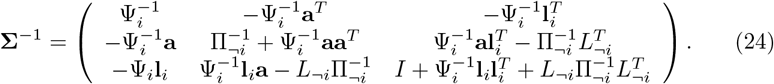

Since we have the complete joint distribution, we can easily extract the marginals of the neural activity by taking the corresponding blocks of the mean and covariance matrices.

#### 5.3.2 Maximum likelihood via expectation-maximization

The CoTuLa model is a latent state model. Thus, parameter inference can be achieved by performing expectation-maximization (EM). In this section, we derive the update rules for EM optimization. To do so, we first must determine the complete log-likelihood. From this, we derive the E-step, followed by the M-step.

##### Complete likelihood

Using the joint distribution calculated above, we can write the log-likelihood over all random variables as

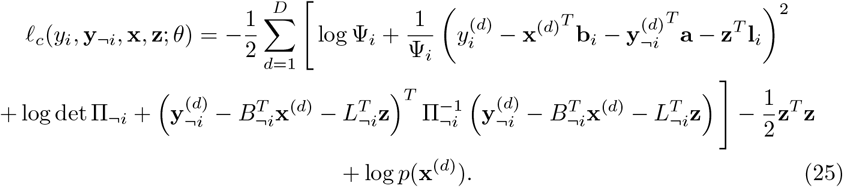

In general, we will ignore the contribution from the density of **x** since it is observed and has no parents in the graphical model.

##### E-step update

To perform the E-step, we need to calculate the averaging distribution *q*(**z**|𝒟 ^(*d*)^; *θ*) with a dataset 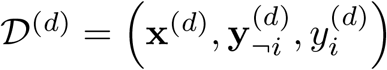. Note that

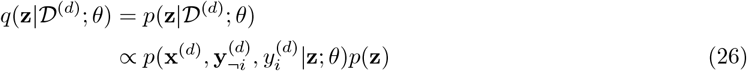

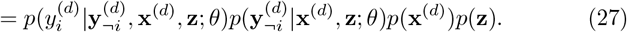

Ultimately, this expression can be written as a Gaussian in **z** with mean ***μ***^(*d*)^ and covariance **Σ**, i.e.

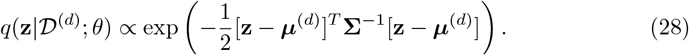

Collecting the quadratic terms gives us the inverse covariance matrix:

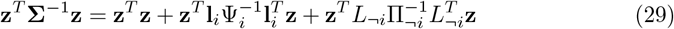

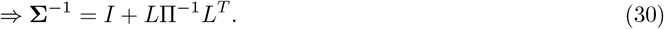

Next, we examine all the linear terms in **z**:

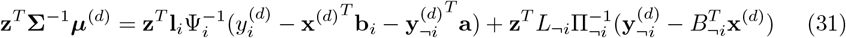

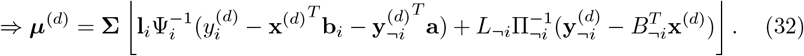

The statistics of the unobserved variables, given by

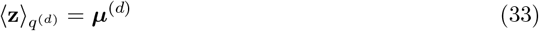

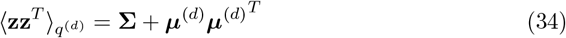

will become relevant in the M-step.

##### M-step update

To calculate the M-step, we take the expectation of the complete log-likelihood over the averaging distribution *q*(**z** | 𝒟). Note that we ignore the prior distribution for **x** as it will not be relevant for any gradients. The expected complete log-likelihood is given by

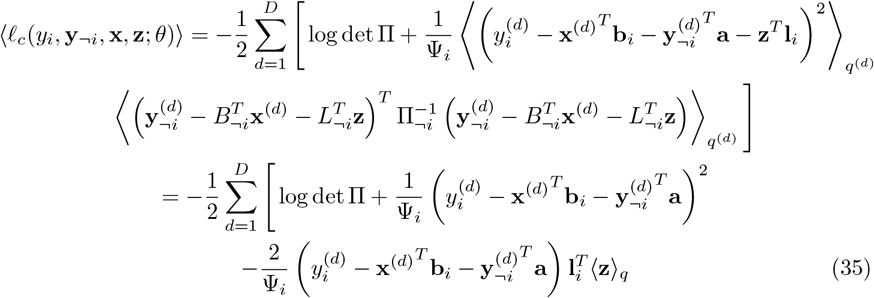

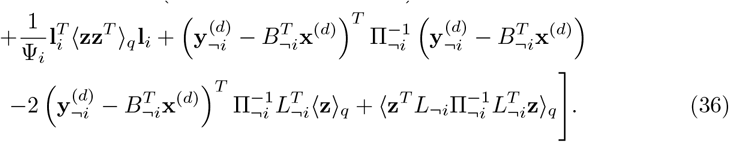

Note that the last expectation can be written as

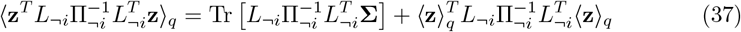

#### 5.3.3 Intercepts and standardizing

In practice, we often want to include an intercept term in our models. With an intercept term, the CoTuLa model becomes:

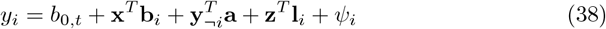

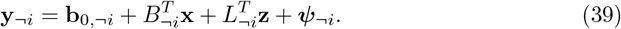

In this case, the marginal distribution (conditioned on the stimulus **x**) of the neural activity is given by

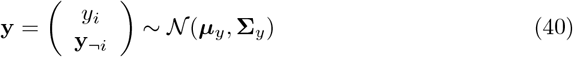

where

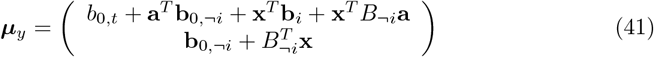

and **Σ**_*y*_ is unchanged. This gives log-likelihood:

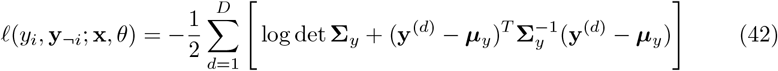

Taking the gradient of the marginal log-likelihood with respect to the intercept terms gives

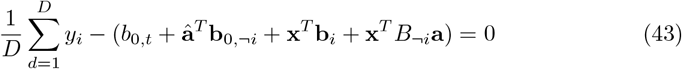

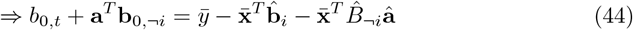

and

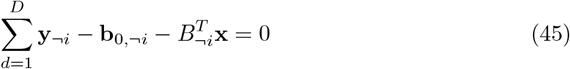

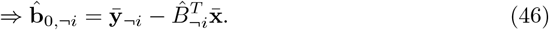

implying that

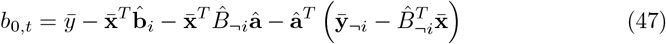

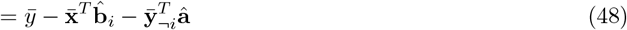

Note that if we center the inputs, i.e. 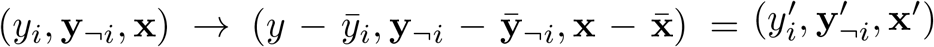, we would find that the intercepts are zero. Thus, performing CoTuLa model inference on the data 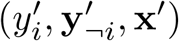 requires that we need not fit an intercept. However, we must transform back to the non-centered space. In such a case, we have

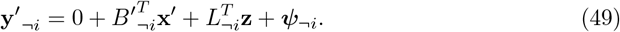

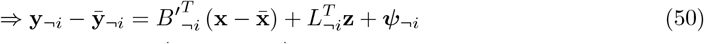

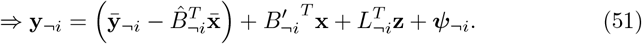

Thus, we have the same intercept formula as above, and 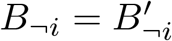. Similarly,

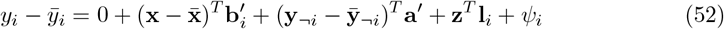

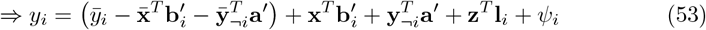

Implying that 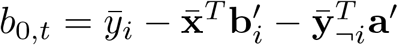, as above.

If we standardize the data in addition to centering, then we have the following:

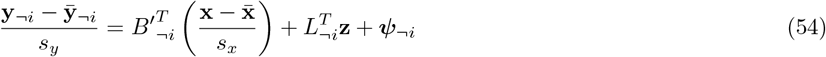

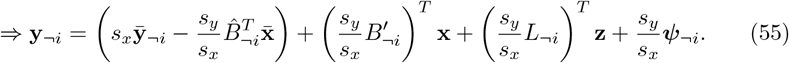

### 5.4 Parametric inference in the dynamical CoTuLa model

#### 5.4.1 OLS biases for the univariate autoregression model

Here we derive the ordinary least-squares regression-based estimates for the coupling and tuning parameters in the model, in the presence of temporally correlated noise. For simplicity, we consider the single-variable model:

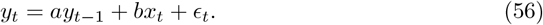

Assuming stationarity, we can only consider the zero-intercept model, by centering the data such that 𝔼 [*x*] = 0 and 𝔼 [*y*] = 0. Note that 𝔼 [·] ≡ 𝔼 [·]_*t*_ indicates an average over time samples, unless otherwise specified. From Eq. 56, we can write

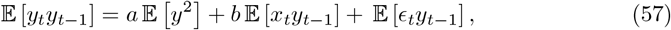

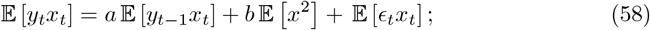

same-time indices are omitted in the averages, such that 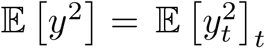. Rewriting, the parameters of the model (*a, b*) have the following relationship with the second-order statistics of the variables:

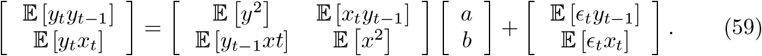

On the other hand, it is straightforward to derive that the OLS estimator for *a* and *b* is given by

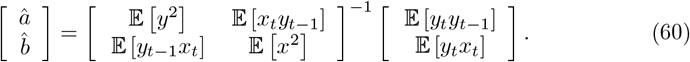

In the limit where the noise *ϵ*_*t*_ is completely uncorrelated with *y*_*t*−1_ or *x*_*t*_, the OLS regression would give exact estimates *â* = *a* and 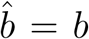. When there are correlations, however, Eq. 59 and Eq. 60 indicate that there are systematic biases:

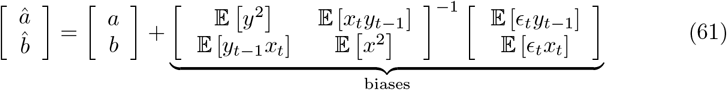

We model the temporally correlated noise using a separate autoregressive process for *z*, and for convenience assume that the external stimulus *x* is also described by an autoregressive model:

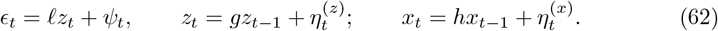

Note that *ϵ*_*t*_ and *y*_*t*−1_ are correlated due to the temporal correlation of *ϵ*_*t*_ and the structure of the dynamical system. (Also see Fig. 5a.) We can write down the correlations under these models,

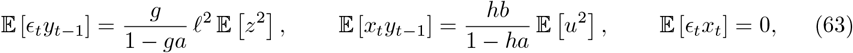

to rewrite the biases in Eq. 61:

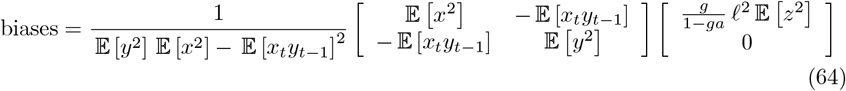

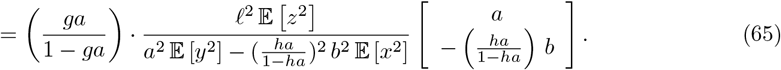

Note how the expression is written in terms of the strengths of the three source terms *ay, bx* and *ℓ z*. Finally, the normalized biases are written as:

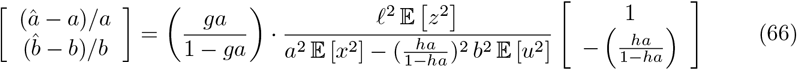

Aside from the three source terms, the normalized biases only depend on *ha* and *ga*, which parameterize the strengths of temporal correlation in the stimuli and the noise, respectively.

#### 5.4.2 Parametric inference by expectation-maximization

We use an Expectation-Maximization (EM) algorithm for inferring the dynamical parameters in the presence of the unobserved source *Z*. Given observed data 𝒟= (*X, Y*), we want to maximize the log-likelihood of observed data, which is lower bounded as:

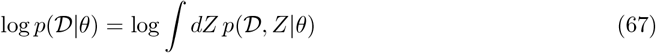

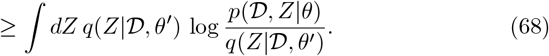

the bound is tight when *q*(*Z* | 𝒟, *θ*) closely approximates *p*(*Z* | 𝒟, *θ*). In our model, the conditional distribution *p*(*Z* | 𝒟, *θ*) can be written as a multivariate normal form (see below for details), so we simply let *q* have the same functional form as *p*(*Z* | 𝒟, *θ*).

The EM inference algorithm iterates between the E-step and the M-step. In the E-step, we construct the *q*(*Z*) distribution by evaluating the conditional distribution *p*(*Z*| 𝒟, *θ*^′^) at the last estimate *θ*^′^. In the M-step, we construct the expected complete-data log-likelihood, *ℓ*_*q*_(*θ*) ≡ 𝔼 [log *p*(𝒟, *Z*|*θ*)]_*q*(*Z*)_. The goal of the M-step is to find the *θ* that maximizes *ℓ*_*q*_(*θ*). Model support can be utilized in the M-step. We derive the full expressions of the complete-data log-likelihood ℒ(*θ*), the conditional distribution *q*(*Z*) = 𝒩 (*Z*|***μ***, Λ), and the expected log-likelihood *ℓ*_*q*_(*θ*) in the following.

##### The complete-data log-likelihood

The joint distribution of the observed data 𝒟 and the unobserved *Z* can be written as

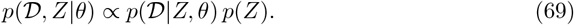

In our current model, each of the two terms in Eq. 69 has a functional form of a normal distribution. The first term (the “likelihood” term) is essentially given by the Gaussian noise assumption:

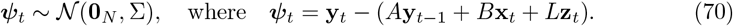

We introduce *trajectory vector notation*, accented by a tilde (∼), which stacks a multivariate time series (an ordered list of vectors) into a single long vector. For example, 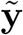 is a *TN* -vector whose *t*-th block is 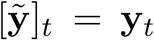. We can similarly define 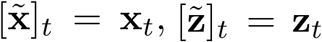 and 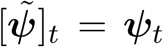 and so on for other time-indexed vectors in the model. We introduce block diagonal matrices 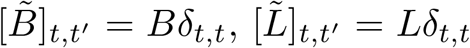, and 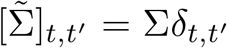, to write 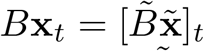 and 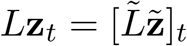. We also construct a *TN* × *TN* matrix 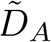, defined block-wise as 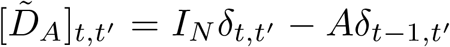, to write 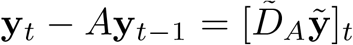. Then Eq. 70 can be written concisely as

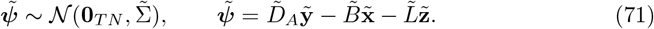

The second term (the “prior” on **z**) is given by the generative model for **z** (Eq. 6):

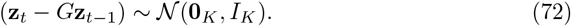

Using the trajectory notation, we introduce a *TK* ×*TK* block matrix 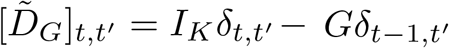, to write 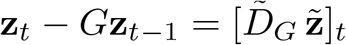. Rewriting,

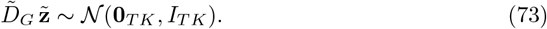

Writing the joint distribution as a function of the parameters *θ* = (*A, B, L*, Σ, *G*), we get the complete data likelihood function. The complete likelihood has a form of a normal distribution in our model, being a product of two normal distributions. Let ℒ (*θ*) be the corresponding log likelihood of having complete data (𝒟, *Z*):

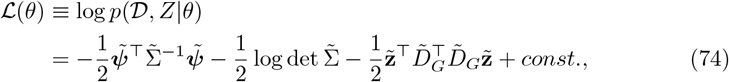

up to terms that are constants of 𝒟, *Z* or *θ*. It is useful to write in a form where the 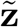 terms are clearly separated. We introduce an auxiliary trajectory vector 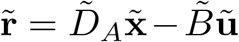, with 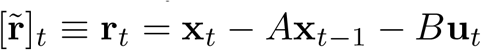. From 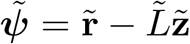, we can write

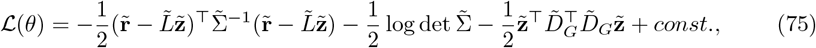

##### The conditional distribution

Because the conditional distribution *p*(*Z*| 𝒟, *θ*) is a normal distribution in the current model, it is fully characterized by an *NK*-dimensional mean vector ***μ*** and an *NK* × *NK* covariance matrix Λ:

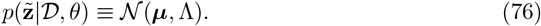

To find ***μ*** and Λ, we expand and collect the 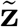-dependent terms in the joint distribution, and match the coefficients:

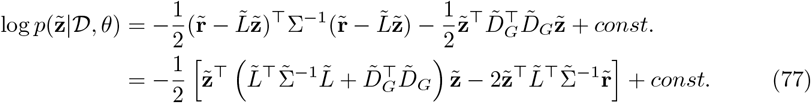

We get

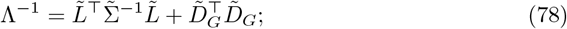

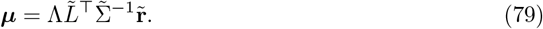

The inverse covariance matrix Λ^−1^ is a symmetric block tridiagonal Toeplitz matrix. Specifically, the (*t, t*^′^) block is written as

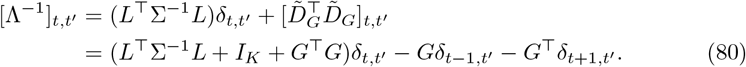

It is possible to iteratively calculate each (*t, t*) block of its inverse matrix, [Λ]_*t,t*_^′^ (see Section 5.4.3). Using [Λ]_*t,t*_^′^, the mean ***μ*** can be calculated block-by-block as

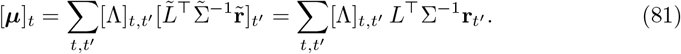

##### The expected log-likelihood

With 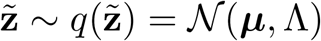, the expected log-likelihood *ℓ*_*q*_(*θ*) can be calculated as:

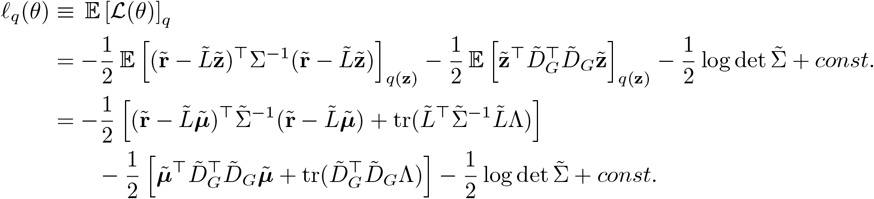

up to terms that are constants of *θ*. We can calculate each term by the *t*-blocks. The quadratic term from the likelihood:

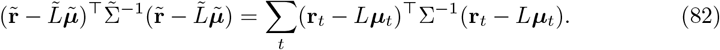

The trace term from the likelihood:

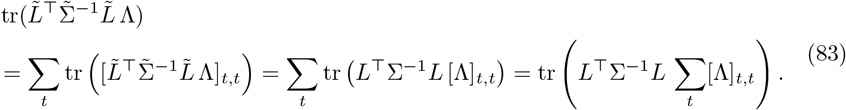

The quadratic term from the prior, from the construction of 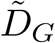:

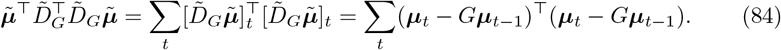

The trace term from the prior:

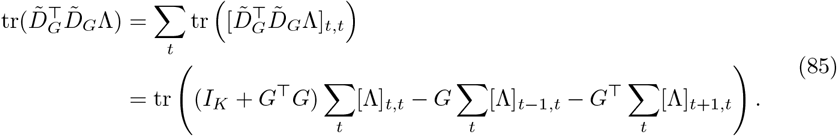

The logdet term:

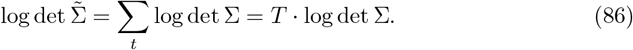

#### 5.4.3 Inversion of a symmetric block-tridiagonal toeplitz matrix

**Problem:** We are interested in the inverse of a symmetric block-tridiagonal Toeplitz matrix of the following form:

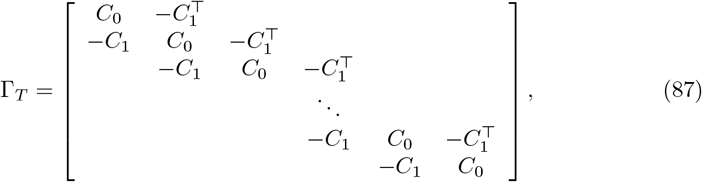

where each block is an *N* × *N* matrix, and there are *T* square blocks on each side (indicated by the subscript in Γ_*T*_). The diagonal block *C*_0_ is a symmetric matrix. Our goal here is to evaluate the inverse matrix 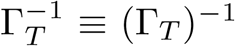 block by block. Specifically, we want to find an expression for the *N* × *N* block of the inverse matrix, 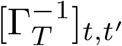, for all 1 ≤ *t, t*^′^ ≤ *T*. Throughout this section, we use the combination of square brackets and subscript for block indexing.

**Solution:** We start by writing down a recursive relationship in the construction of the block matrix. When there is only 1 block, clearly Γ_1_ = *C*_0_ and 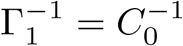. For all *k >* 1,

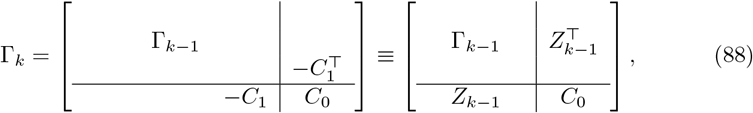

where *Z*_*k*−1_ ≡ [0_*N*×*N*_, …, −*C*_1_] is a row of *k* −1 blocks, with the last block [*Z*_*k*−1_]_*k*−1_ = −*C*_1_ and otherwise zero. The dividing lines and the uneven alignment are just to visually emphasize the different sizes of the sub-matrices; we drop these styles below.

Inversion of the block matrix (Eq. 88) gives

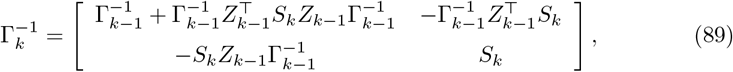

where 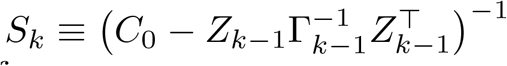. From the block-sparse structure of *Z*_*k*−1_, we can simplify

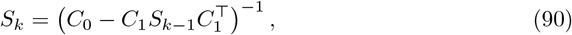

where 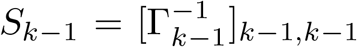 by construction. Similarly, if we define *Y*_*k*_ to be the last block-row of 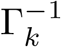 for all *k*, we can also simplify 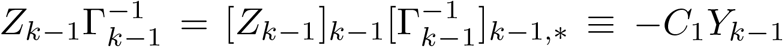. Then we rewrite:

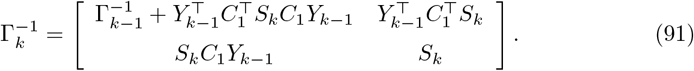

The bottom row of Eq. 91 gives a recursive relationship for *Y*_*k*_ = [*S*_*k*_*C*_1_*Y*_*k*−1_, *S*_*k*_]. Specifically, we note that 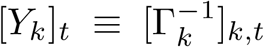 has a chain-like form. When *t* = *k*, we have [*Y*_*k*_]_*k*_ = *S*_*k*_ by construction. When *t < k*,

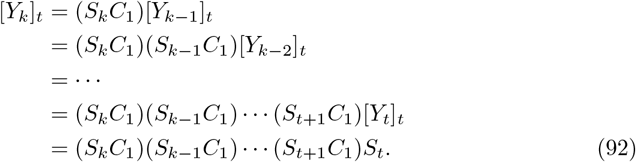

We can pre-compute the sequence of *S*_*k*_’s for all 1 ≤ *k* ≤ *T*, starting from 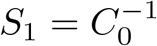, and proceeding according to Eq. 90. Then we are ready to calculate 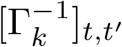. Due to symmetry, it is sufficient to only consider blocks with *t* ≥ *t*^′^. When *t*^′^ ≤ *t* = *T*, we have 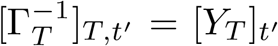, by definition of *Y*. When *t*^′^ ≤ *t < T*, using the upper left block of Eq. 91:

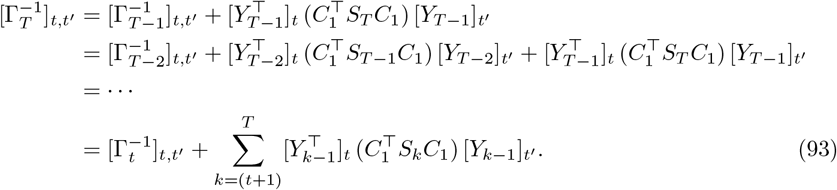

In fact, the *t*^′^ ≤ *t* = *T* solution is a special case of the last expression, because 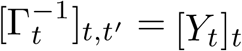 and the summation range for the second term is empty when *t* = *T*.

Putting together, for all *t*^′^ ≤ *t* ≤ *k*,

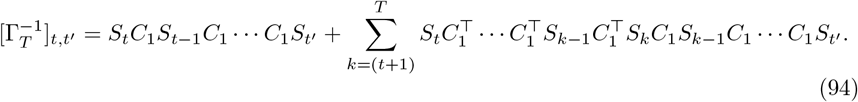

### 5.5 Selection procedures

#### 5.5.1 Union of Intersections (UoI)

We performed sparse coupling and tuning fits on the target neuron (separately for coupling and tuning), using UoI_Lasso_, a recent method that has been demonstrated to achieve improved inference in systems neuroscience models [18, 19]. We then determined the model supports by taking the parameters that were non-zero in the UoI_Lasso_ solution.

#### 5.5.2 Normalization-and-cutoff

We first performed separate coupling and tuning fits using OLS estimates. We normalized each of the resulting coupling and tuning matrices by the rows and columns, then applied a cutoff to the elements of each normalized matrix to determine the model supports. The row/column normalization was motivated by the column-wise correlation structure in the OLS estimate errors when data was generated from a CoTuLa model, in the univariate case shown in Section 5.4.1. The value of the cutoff was optimized by the Bayesian Information Criterion.

### 5.6 Analysis of estimated parameters

#### Normalized biases

In the synthetic experiment, it is possible to calculate the biases of estimated parameters because we have access to the ground truth values. The normalized bias of a parameter estimate was calculated element-wise, (*v*_est_ − *v*_true_)*/* | *v*_true_, where *v* is each element of the parameter matrix. We averaged over all elements where the ground truth value is non-zero (|*v*| *>* 0).

#### Tuning and coupling strengths

For tuning, we calculated a tuning modulation strength for each neuron. Tuning modulation was defined as the norm of the *M* = 2 tuning parameters (visual cortex data), the minimum to maximum distance of the tuning curves generated across frequencies using the Gaussian basis functions (auditory cortex data), and the peak height of the tuning curve (hippocampus data). For coupling, in all datasets, we showed the raw coupling parameters in the model supports, and the absolute value of those parameters for summary statistics.

#### Fraction of Variance for the Dynamic CoTuLa parameters

The relative coupling contribution is calculated as the variance fraction, (∑_*t*_ ‖*Â***y**_*t*−1_‖^2^)*/*(∑_*t*_ ‖**y**_*t*_‖^2^) (see Eq. 5), where the sums were taken over all time points included in each fit, and *Â* is the estimate from this fit. The tuning, latent, and error contributions were similarly calculated using 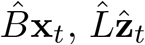 and 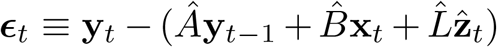 in the numerator, respectively. Note that the variance fractions do not necessarily sum to 1 because the terms in the model are not all orthogonal.

#### Box plot

Circles mark the median value. Bars extend from *Q*_1_ to *Q*_3_, where *Q*_1_ and *Q*_3_ are the first and the third quartiles (25% and 75%) of the distribution. This is the standard box plot without the whiskers.

#### Statistical tests

We used Wilcoxon signed-rank tests/Wilcoxon rank-sum tests, and used the following markers to indicate the significance levels in the plot: *** (*p* ≤ 0.001), ** (*p* ≤ 0.01), * (*p* ≤ 0.05), and no marker (*p >* 0.05).

### 5.7 Structural non-identifiability in the static CoTuLa model

In this section, we show that the static CoTuLa model is structurally non-identifiable. Furthermore, we prove that sufficient sparsity in the static CoTuLa model remedies the non-identifiability.

#### 5.7.1 Deriving the identifiability subspace

Recall that the marginal log-likelihood is given by

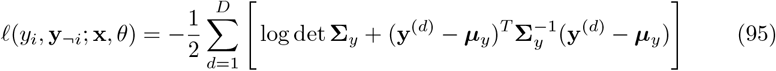

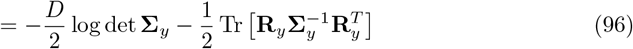

where

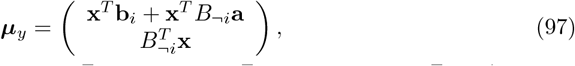

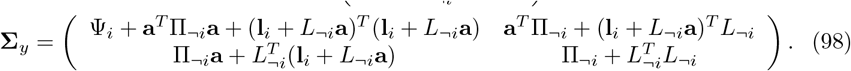

Here, we consider an identifiability issue in the “separable” sense, i.e. offsets that can be applied to the parameters such that the mean ***μ***_*y*_ and covariance **Σ**_*y*_ are separately unchanged. If these quantities remain unchanged, the log-likelihood will necessarily be unchanged. The “separable” case is in contrast to the scenario in which we change the parameters such that the log-likelihood is unchanged, but through its overall computation (rather than due to the mean and covariance remaining unchanged).

We apply offsets to **l**_*i*_, Ψ_*i*_, **a**, and **b**_*i*_. Specifically, suppose we apply some offset ***δ*** to **l**_*i*_:

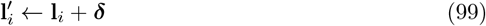

and define the quantity

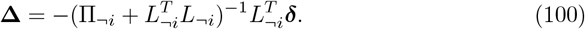

Then, we apply the following transformations to Ψ_*i*_, **a** and **b**_*i*_:

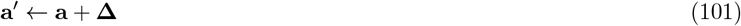

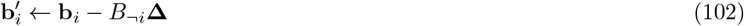

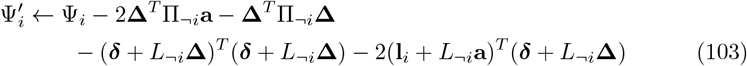

The bottom block of the mean is necessarily unchanged, since we have not modified *B*_¬*i*_. Meanwhile, the top block, with the new parameter configuration, becomes

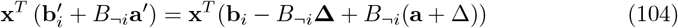

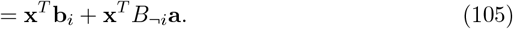

Thus, ***μ***_*y*_ is unchanged with the new parameter configuration. In the covariance matrix **Σ**_*y*_, the bottom right quadrant is necessarily unchanged, since we do not modify any of the constituent parameters. The bottom left component (which is identical in content to the top right component), under the new configuration, is given by

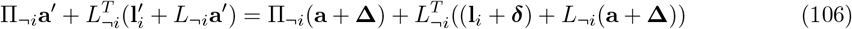

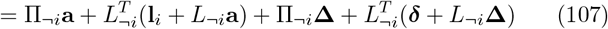

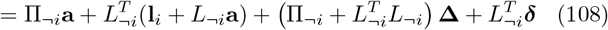

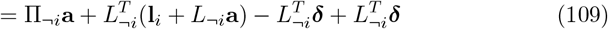

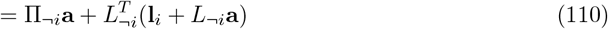

and so is unchanged. Lastly, we consider the top-left component of the covariance matrix:

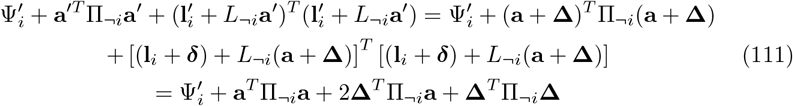

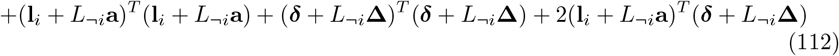

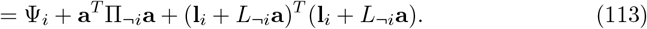

Thus, the variance for the target neuron is unchanged. These offsets specify a family of solutions, given a *B*_¬*i*_, *L*_¬*i*_, and Π_¬*i*_. Importantly, however, this family is restricted to where Ψ_*i*_ is positive.

#### 5.7.2 Sufficient model sparsity removes non-identifiability

*Proof*. The only free parameters lie in the *K*-dimensional subspace determined by ***δ***. We can rewrite the identifiability transform equations as

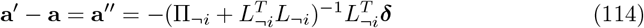

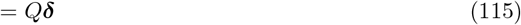

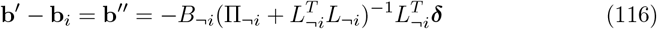

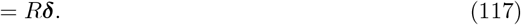

In these equations, the subset of **a**^′ ′^ and **b**^′ ′^ which start at 0 due to the model support must remain at 0 to preserve the model support. Thus, we are only concerned with the subset of the parameters that are constrained to be 0. Call this subset (of rows)

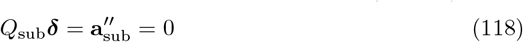

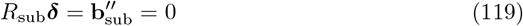

where *Q*_sub_ and *R*_sub_ are the *Nk*_*C*_ × *K* and *Mk*_*T*_ × *K* linear transforms. This linear system can collectively be written as the (*Nk*_*C*_ + *Mk*_*T*_) × *K* linear system

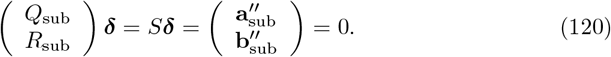

When *K* ≤ *Nk*_*C*_ + *Mk*_*T*_ and *S* is full-rank, the only solution is ***δ*** = 0.

Otherwise, there will be a family of non-trivial solutions which live in the kernel of *S*. So, solutions with higher sparsity permit a higher latent dimensionality which still having a unique support-preserving solution. Thus, for a sufficiently sparse estimate with low enough latent dimensionality, a check of the rank of a matrix is sufficient to determine whether the identifiability subspace has been constrained through fixing the support. Note that this determines conditions on identifiability for both ground-truth parameters or estimated parameters in the static CoTuLa model.

## 6 Author Contributions

Conceptualization: KEB Project Management: KEB

Funding acquisition: KEB, CK, LF

Methodology: PS, JL, JHB, SB, KEB Software: PS, JL, JHB

Visualization: PS, JHB, KEB

Writing – original draft: PS, JHB, SB, KEB

Writing – review and editing: PS, JHB, JL, CK, LF, SB, KEB,

## 7 Acknowledgements

K.E.B. was funded by a Lawerence Berkeley National Laboratory LDRD for the Neural Systems and Data Science Lab, National Institute of Neurological Disorders and Stroke Grant RNS118648A; P.S. was funded by a DOD NDSEG; J.H.B. was funded by a UCSF Weill Neuroscience Institute (awarded to L.F., C.K., and K.E.B.). J.H.B. and K.E.B. thank Asadali Abbas Hazariwala for assistance with computational studies.

## Notes

### Competing Interest Statement

The authors have declared no competing interest.

