## Supplementary Material for "Resolving Non-identifiability Mitigates Bias in Models of Neural Tuning and Functional Coupling"

### Supplementary Information: Resolving Non-identifiability Mitigates Biases in Models of Neural Tuning and Functional Coupling

#### 1 Alternative Inference Procedures

The structural non-identifiability elevates the issue of selection in parameter inference for the static CoTuLa model. Specifically, a sufficiently sparse number of non-zero tuning and coupling parameters must be identified to ensure that their values can be accurately estimated, thereby avoiding the simultaneous equations bias. Thus, parameter inference in the static CoTuLa model can be modularized into a selection procedure, which identifies the non-zero parameters, and an estimation procedure, which estimates their values given a model support. In this section, we detail alternative selection procedures we considered.

**L1-Penalized Sparse CoTuLa Inference.** One natural approach to performing selection in the static CoTuLa model is to apply an $\ell_1$ penalty to the tuning and coupling parameters during expectation-maximization. However, since these parameters may have different sparsity levels, a different $\ell_1$ penalty must be applied to each set. In practice, this would occur during the M-step of the EM optimization when maximizing the expected complete log-likelihood. Thus, the M-step would consist of minimizing the expression

$$\ell_M(\theta;\mathcal{D}) = -\langle \ell_c(\theta;\mathcal{D})\rangle + \lambda_1|\mathbf{a}|_1 + \lambda_2|\mathbf{b}_i|_1 \tag{1}$$

where we assume that the tuning penalties are applied equally across both target and non-target parameters. Such an optimization would require cross-validating over a grid of $(\lambda_1, \lambda_2)$ combinations.

In practice, only one $\lambda$ penalty can be used at a time. We can sidestep this issue by rescaling the parameters during optimization. Let $r = \lambda_2/\lambda_1$ and

$$\mathbf{b}_i' = r\mathbf{b}_i \tag{2}$$

so that

$$\begin{aligned}
\ell_M(\mathbf{a},\mathbf{b}_i,\theta;\mathcal{D}) &= -\langle \ell_c(\mathbf{a},\mathbf{b}_i,\theta;\mathcal{D})\rangle + \lambda_1|\mathbf{a}|_1 + \lambda_2|\mathbf{b}_i|_1 &(3)\\
&= -\langle \ell_c(\mathbf{a},\frac{1}{r}\mathbf{b}_i',\theta;\mathcal{D})\rangle + \lambda_1|\mathbf{a}|_1 + \frac{\lambda_2}{r}|\mathbf{b}_i'|_1 &(4)\\
&= -\langle \ell_c(\mathbf{a},\frac{1}{r}\mathbf{b}_i',\theta;\mathcal{D})\rangle + \lambda_1|\mathbf{a}|_1 + \lambda_1|\mathbf{b}_i'|_1 &(5)\\
&= \ell_M'(\mathbf{a},\mathbf{b}_i',\theta;\mathcal{D}). &(6)
\end{aligned}$$

The expressions $\ell_M$ and $\ell_M'$ are equivalent aside from a reparameterization. Thus, the minimization we want to achieve,

$$\mathbf{a}^*, \mathbf{b}_i^* = \underset{\mathbf{a},\mathbf{b}_i}{\arg\min}\, \ell_M(\mathbf{a},\mathbf{b}_i,\theta;\mathcal{D}) \tag{7}$$

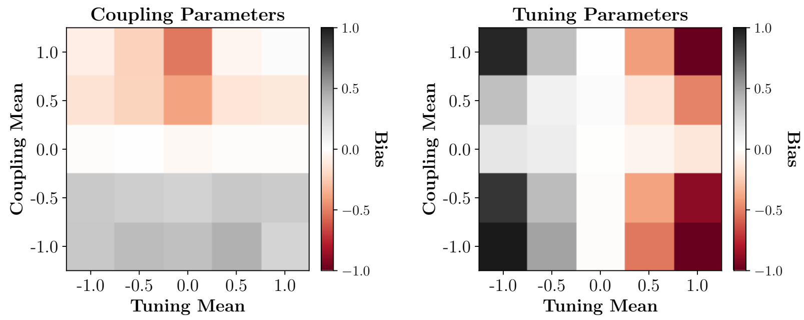


**Figure S.1: L1-penalized sparse** Plotted are the biases for the the CoTuLa model with inference performed with $L_1$ regularized expectation maximization. As in Figure 3, we plot bias (given by color bars) for the tuning and coupling parameters separately as a function of the means of the distributions from which the coupling and tuning parameters were drawn.

can be achieved by instead minimizing

$$\mathbf{a}^*, \mathbf{b}_i'^* = \underset{\mathbf{a}, \mathbf{b}_i'}{\arg\min}\, \ell_M'(\mathbf{a}, \mathbf{b}_i', \theta; \mathcal{D}). \tag{8}$$

However, we want the solution that minimizes $\ell_M$, so after performing the optimization, we need to transform back to the desirable parameters:

$$\mathbf{b}_i^* = \mathbf{b}_i'^*/r. \tag{9}$$

**L1-Penalized Sparse CoTu Inference.** If we enforce sparsity in this fashion via the static CoTuLa model, the most natural comparison to the tuning and coupling model is via a similar sparse optimizer. Specifically, the loss function would be simply be the mean-squared error, with the additional penalties:

$$\ell_{TC}(\mathbf{a}, \mathbf{b}_i; \mathcal{D}) = \sum_{d=1}^{D} \left( y^{(d)} - {\mathbf{x}^{(d)}}^T \mathbf{b}_i - {\mathbf{y}_{\neg i}^{(d)}}^T \mathbf{a} \right)^2 + \lambda_1 |\mathbf{a}|_1 + \lambda_2 |\mathbf{b}_i|_1 \tag{10}$$

This optimization problem is ultimately a linear regression with lasso penalty, with some parameters penalized differently than others. Thus, it can easily be solved using a cross-validation grid to determine the best $(\lambda_1, \lambda_2)$ configuration. In Fig. S.1 we display CoTuLa parameter bias for coupling (left) and tuning (right) parameters as a function of means of the distributions from which the parameters were drawn. Here, we see that bias persists and is substantially larger than with the two-stage procedure utilized in Figure 3e-h, and is qualitatively similar in magnitude to the results for CoTuLa inference with no selection (Figure 3j-l). We note that, in contrast to the resuls seen in Figure 3, we observe negative coupling biases. This likely arises due to the fact that the $\ell_1$-penalized EM solver is prone to false negatives, which will automatically produce parameter estimates with negative coupling bias.

### 2 Selection Accuracy of UoI

In Fig. S.2, we present results demonstrating the accuracy of variable selection (i.e., inference of support) using the Union of Intersections method (UoI) as a function of the means of the coupling and tuning parameter distributions. Selection accuracy is

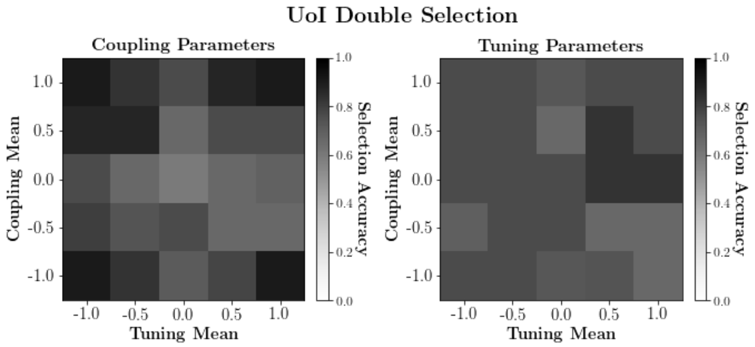


**Figure S.2:** Results demonstrating the accuracy of variable selection (i.e., inference of support) using the Union of Intersections method (UoI). The color bar indicates the selection accuracy for coupling (left) and tuning (right) parameters as a function of the means of the coupling and tuning parameter distributions.

quantified as the Jacard index of the inferred and actual supports, where 0 corresponds to no overlap between the inferred support set and the ground truth, and 1 corresponds to perfect recovery of the support. As we have demonstrated in many other papers [1–4], UoI excels at feature selection for both tuning and coupling. We note that support recovery for the coupling terms is particularly challenging given the latent influences. Despite this challenging situation, UoI is able to obtain near-perfect coupling selection accuracy in conditions where the means of the underlying parameters are large (e.g., corners of the matrix.), as these are the regions of the parameter space that impart the largest signal in the modeled responses. The high degree of overlap in the data generation process for tuning curves explains the reduced performance in that case (there is a very high degree of multicollinearity amongst the tuning parameters.) Prior work applying UoI to real neural data for tuning curve estimation [2] indicates that this is not an issue in real data.

#### 2.1 Bias in Poisson CoTuLa Models

We devised a Poisson CoTuLa model in order to examine whether the omitted variables are present in data generated from the model. The model is structured similarly to that of the linear-Gaussian instantiation, except it replaces the addition of private Gaussian variability with a Poisson distribution. Thus, the model further requires an exponential nonlinearity to ensure non-negative means are passed to the Poisson distribution. We chose this model based on Poisson factor analysis [5]. The Poisson triangular model is given by

$$\mu_{\neg i} = \exp\left(\mathbf{b}_{\neg i}^T\mathbf{x} + \mathbf{l}_{\neg i}^T\mathbf{z}\right) \tag{11}$$

$$y_j = \text{Poisson}(\mu_{\neg i} \cdot \Delta) \tag{12}$$

$$\mu_t = \exp\left(\mathbf{b}_i^T\mathbf{x} + \mathbf{y}_{\neg i}^T\mathbf{a} + \mathbf{z}^T\mathbf{l}_i\right) \tag{13}$$

$$y_i = \text{Poisson}(\mu_i \cdot \Delta). \tag{14}$$

We use $\neg i$ to index over the non-target neurons. We emphasize that the Poisson distribution is applied independently to each neuron (rather than applied in a multivariate fashion). This implies that $\mathbf{y}_{\neg i} = [y_j]_{j\neq t}$. Furthermore, $\Delta$ denotes the bin width. We

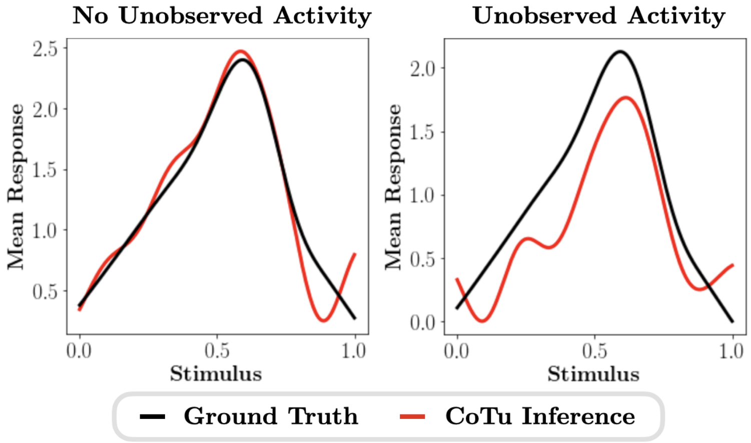


**Figure S.3: Omitted variables biases are qualitatively similar when the neural response is modeled as a Poisson distribution.** Ground truth tuning curves (black) and CoTu model tuning curves (red) inferred from data generated from a Poisson CoTuLa model with no unobserved activity (left) and unobserved activity (right). Tuning curves depict the mean response (after the exponential nonlinearity) as a function of the stimulus.

can write this model for $D$ samples as

$$\begin{aligned}
\boldsymbol{\mu}_{\neg i} &= \exp\left(\mathbf{X}\mathbf{B}_{\neg i} + \mathbf{Z}\mathbf{L}_{\neg i}\right) && (15)\\
\mathbf{Y}_{\neg i} &= \text{Poisson}\left(\boldsymbol{\mu}_{\neg i} \cdot \Delta\right) && (16)\\
\boldsymbol{\mu}_{i} &= \exp\left(\mathbf{X}\mathbf{b}_{i} + \mathbf{Y}_{\neg i}\mathbf{a} + \mathbf{Z}\mathbf{l}_{i}\right) && (17)\\
\mathbf{y}_{i} &= \text{Poisson}\left(\boldsymbol{\mu}_{i} \cdot \Delta\right) && (18)
\end{aligned}$$

where, importantly, the exponential nonlinearity and Poisson distribution are applied element-wise.

We generated data from the Poisson CoTuLa model with tuning parameters chosen with Gaussian basis functions and coupling parameters chosen from a Gaussian distribution centered at zero. We generated datasets from a CoTuLa model with no unobserved activity (i.e., latent factors all set to zero) and one with unobserved activity (i.e., positive noise correlations). From the data, we fit a Poisson CoTu model (i.e., a generalized linear model with exponential link function) to the data. We found that, when no unobserved activity was present in the data, the model faithfully captured the tuning parameters (Fig. S.3, left). Meanwhile, if the unobserved activity was present in the data, the CoTu tuning curve underestimated the ground truth tuning curve (Fig. S.3, right). Together, these results demonstrate that the omitted variable bias is present in Poisson CoTu models, with a qualitatively similar structure to the biases observed in the linear-Gaussian CoTuLa models.

### 3 OLS biases for the univariate static CoTuLa model

For a univariate static CoTuLa model, we can derive the OLS biases when fitting a CoTu model. Unfortunately, the univariate model is not identifiable and the biases will depend on parameters impacted by the identifiability. The univariate static CoTuLa model is

$$y_i = b_i x + a y_{\neg i} + l_i z + \psi_i \tag{19}$$
$$y_{\neg i} = b_{\neg i} x + l_{\neg i} z + \psi_{\neg i}. \tag{20}$$

We will assume that the variables ($x$, $z$, $\psi_i$, $\psi_{\neg i}$) are mean-centered and uncorrelated (note Eqs 19 and 20 can induce correlations between $x$, $y_i$, etc.). The OLS estimates for $a$ and $b_i$ in the CoTu model are

$$\begin{bmatrix}\hat{a}\\ \hat{b}_i\end{bmatrix} = \begin{bmatrix}\langle y_{\neg i}^2\rangle & \langle y_{\neg i}x\rangle\\ \langle y_{\neg i}x\rangle & \langle x^2\rangle\end{bmatrix}^{-1}\begin{bmatrix}\langle y_i y_{\neg i}\rangle\\ \langle y_i x\rangle\end{bmatrix} \tag{21}$$

From Eq 19, we can derive

$$\langle y_i y_{\neg i}\rangle = b_i\langle y_{\neg i}x\rangle + a\langle y_{\neg i}^2\rangle + l_i\langle y_{\neg i}z\rangle + \langle y_{\neg i}\psi_i\rangle \tag{22}$$
$$\langle y_i x\rangle = b_i\langle x^2\rangle + a\langle y_{\neg i}x\rangle + l_i\langle xz\rangle + \langle x\psi_i\rangle \tag{23}$$

which can be rearranged as

$$\begin{bmatrix}a\\ b_i\end{bmatrix} = \begin{bmatrix}\langle y_{\neg i}^2\rangle & \langle y_{\neg i}x\rangle\\ \langle y_{\neg i}x\rangle & \langle x^2\rangle\end{bmatrix}^{-1}\left(\begin{bmatrix}\langle y_i y_{\neg i}\rangle\\ \langle y_i x\rangle\end{bmatrix} - l_i\begin{bmatrix}\langle y_{\neg i}z\rangle\\ \langle xz\rangle\end{bmatrix} - \begin{bmatrix}\langle y_{\neg i}\psi_i\rangle\\ \langle x\psi_i\rangle\end{bmatrix}\right) \tag{24}$$

and simplified to

$$\begin{bmatrix}a\\ b_i\end{bmatrix} = \begin{bmatrix}\langle y_{\neg i}^2\rangle & \langle y_{\neg i}x\rangle\\ \langle y_{\neg i}x\rangle & \langle x^2\rangle\end{bmatrix}^{-1}\left(\begin{bmatrix}\langle y_i y_{\neg i}\rangle\\ \langle y_i x\rangle\end{bmatrix} - l_i\begin{bmatrix}\langle y_{\neg i}z\rangle\\ 0\end{bmatrix}\right). \tag{25}$$

Plugging this into the equation for the OLS estimates gives

$$\begin{aligned}\begin{bmatrix}\hat{a}\\ \hat{b}_i\end{bmatrix} &= \begin{bmatrix}a\\ b_i\end{bmatrix} + \begin{bmatrix}\langle y_{\neg i}^2\rangle & \langle y_{\neg i}x\rangle\\ \langle y_{\neg i}x\rangle & \langle x^2\rangle\end{bmatrix}^{-1} l_i\begin{bmatrix}\langle y_{\neg i}z\rangle\\ 0\end{bmatrix}\\ &= \begin{bmatrix}a\\ b_i\end{bmatrix} + \frac{l_i\langle y_{\neg i}x\rangle}{\langle y_{\neg i}^2\rangle\langle x^2\rangle - \langle y_{\neg i}x\rangle^2}\begin{bmatrix}\langle x^2\rangle\\ -\langle y_{\neg i}x\rangle\end{bmatrix}.\end{aligned} \tag{26}$$

We can now substitute several expectations derived from Eq 20

$$\begin{aligned}\begin{bmatrix}\hat{a}\\ \hat{b}_i\end{bmatrix} &= \begin{bmatrix}a\\ b_i\end{bmatrix} + \frac{l_i b_{\neg i}\langle x^2\rangle}{(b_{\neg i}^2\langle x^2\rangle + l_{\neg i}^2 + \langle\psi_{\neg i}^2\rangle)\langle x^2\rangle - b_{\neg i}^2\langle x^2\rangle^2}\begin{bmatrix}\langle x^2\rangle\\ -b_{\neg i}\langle x^2\rangle\end{bmatrix}\\ &= \begin{bmatrix}a\\ b_i\end{bmatrix} + \frac{l_i b_{\neg i}\langle x^2\rangle}{l_{\neg i}^2 + \langle\psi_{\neg i}^2\rangle}\begin{bmatrix}1\\ -b_{\neg i}\end{bmatrix}\end{aligned} \tag{27}$$

and get normalized biases

$$\begin{bmatrix}(\hat{a} - a)/|a|\\ (\hat{b}_i - b)/|b|\end{bmatrix} = \frac{l_i\langle x^2\rangle}{l_{\neg i}^2 + \langle\psi_{\neg i}^2\rangle}\begin{bmatrix}b_{\neg i}/|a|\\ -b_{\neg i}^2/|b|\end{bmatrix} \tag{28}$$

The overall scale of the normalized bias depends on $\frac{l_i\langle x^2\rangle}{l_{\neg i}^2 + \langle\psi_{\neg i}^2\rangle}$ which is larger if either the stimuli are large amplitude, the impact of the shared variability on the target-neuron is large, or if the shared or private noise in the non-target neurons is large. Unfortunately, the non-identifiability of the univariate static CoTuLa model means that it is often possible to have $l_i = 0$ (and in general, $l_i$ can undergo an identifiability transform) in the identifiability subspace, which means that zero-bias TC fits are often possible. Multivariate, sparse identifiable biases could be derived, but they will not have nice analytic forms (they will involve matrix inverses).

The coupling additionally bias scales with the ratio of the non-target tuning to the coupling. The tuning bias additionally scales negatively with the ratio of the square of the non-target tuning to the target tuning.
